# Impaired Chylomicron Secretion Results in Reduced Body Fat and Increased Intestinal Fatty Acid Oxidation by Activation of Autophagy

**DOI:** 10.1101/2025.11.19.689245

**Authors:** Khaga R. Neupane, Alexander Karakashian, Maura Mobilia, Olivia Hage, Patrick Tso, Min Liu, Laurent Vergnes, Karen Reue, Scott M. Gordon

**Affiliations:** Saha Cardiovascular Research Center, University of Kentucky, Lexington, KY; Department of Physiology, University of Kentucky, Lexington, KY; Department of Pathology and Laboratory Medicine, Reading Campus, University of Cincinnati, 2180 E Galbraith Road, Cincinnati, OH 45237, USA; Department of Human Genetics, David Geffen School of Medicine, University of California-Los Angeles, Los Angeles, California USA; Department of Medicine, and Molecular Biology Institute, University of California-Los Angeles, Los Angeles, California USA

**Keywords:** chylomicron, body composition, dietary lipid, intestine, enterocyte, DENND5B

## Abstract

Intestinal absorption of dietary lipid is essential for systemic lipid homeostasis; however, elevated postprandial plasma lipid levels are associated with obesity and increased risk for atherosclerotic cardiovascular disease. In humans and rodents, biological sex impacts dietary triglyceride absorption and males tend to have higher postprandial triglyceride levels compared to females, but the physiological basis for this is not well understood. Here, we show that the gene *DENND5B* is associated with body composition in humans and mice and that genetic deletion of *Dennd5b* in mice prevents postprandial plasma triglyceride elevations in both sexes. Our findings establish a role for this protein in intestinal chylomicron secretion and reveal a biological sex-biased differential impact of its deletion on body composition, dietary fatty acid uptake, and intestinal metabolism. Mechanistically, our findings implicate autophagy and mitochondrial beta oxidation in coping with enterocyte lipid accumulation due to impaired chylomicron secretion. This work extends the emerging concept that the intestinal epithelium may play a prominent role in oxidation of diet-derived fatty acids and suggests that this process may contribute to biological sex-based differences in postprandial lipemia and body composition.

**Highlights:**

- The *DENND5B* gene is required for dietary triglyceride absorption and is associated with metabolic phenotypes in humans and mice.
- Disruption of chylomicron secretion differentially affects body composition, dietary lipid absorption, and intestinal metabolic activity in male and female mice.
- In mice, genetic disruption of *Dennd5b* results in lower body fat in both sexes, and increased lean mass only in males.
- Metabolic phenotypes in *Dennd5b*-deficient mice are driven by reduced absorption of dietary lipid characterized by enterocyte lipid retention and increased fecal lipid excretion.
- Impaired chylomicron secretion in *Dennd5b^−/−^* mice induces autophagy-mediated disposal of cellular triglyceride and increased fatty acid oxidation in intestinal tissue.

---

Keywords: chylomicron, body composition, dietary lipid, intestine, enterocyte, DENND5B

## Introduction

Absorption of dietary lipid by the intestine is essential for maintenance of systemic lipid homeostasis ^1^. After consuming a meal containing fat, absorptive enterocytes in the intestinal epithelium rapidly take up ingested fatty acids, re-esterify them to triacylglyceride (TAG), and load the TAG into chylomicrons for secretion to the mesenteric lymph. The mesenteric lymph then delivers TAG-loaded chylomicrons to the circulation for systemic distribution. Postprandial plasma triglyceride levels are associated with body composition and are an independent predictor of cardiovascular risk ^2-5^. Males experience higher postprandial triglyceride levels than females ^6,7^. While the mechanism behind this observation is not well understood, it likely involves a combination of factors including sex differences in intestinal absorption and rates of peripheral lipolysis and uptake of fatty acids.

In addition to acute effects on postprandial plasma TAG levels, intestinal processing of dietary lipids during absorption may impact body composition by influencing the total mass of ingested TAG that ultimately enters the body. Several studies have examined the impact of dietary fat on intestinal metabolism ^8-11^, but few have directly compared males and females. Post-absorption, males and females utilize absorbed TAG differently ^12^. Males tend to channel more ingested fatty acids to beta oxidation in peripheral tissues for immediate energy availability, while females store more TAG in adipose tissue to maintain energy reserves ^13,14^. This likely contributes to the well characterized sex differences in body composition and body fat distribution ^15^. Sex differences in mitochondrial function have been described that may impact fatty acid utilization in specific tissues ^13^. A recent study by Moschandrea et al. demonstrated that disruption of mitochondrial function in the intestinal epithelium impairs chylomicron secretion ^16^. These findings suggest that mitochondrial function is required for the proper trafficking of triglyceride-loaded chylomicrons through the secretory pathway. The impact of biological sex on mitochondrial function in the intestinal epithelium and its potential impact on the absorption of dietary TAG is not well understood.

Impaired absorption of dietary lipids occurs in humans with genetic mutations in genes involved in intracellular TAG reesterification (*MGAT*, *DGAT*), chylomicron formation (*APOB*, *MTTP*), or ER to Golgi trafficking of chylomicrons (*SAR1B*). When this lipid absorption pathway is disrupted, patients experience failure to thrive, steatorrhea after consumption of lipid, and fat-soluble vitamin deficiency ^17-19^. We recently discovered a new component of this pathway, *DENND5B*, which appears to play a role in chylomicron secretion that is downstream of previously described proteins, acting in the post-Golgi trafficking of chylomicrons for secretion from the enterocyte ^20^. Detailed electron microscopy analysis of intestinal tissue demonstrated that formation of chylomicrons in the endoplasmic reticulum, pre-chylomicron transport vesicle trafficking between the ER and Golgi, and the presence of intra-Golgi chylomicrons were intact in *Dennd5b^−/−^* mice. However, we observed an accumulation of chylomicron secretory vesicles (membrane bound vesicles containing multiple chylomicron-sized lipoproteins) within enterocytes and a prominent reduction of extracellular chylomicrons in the lamina propria after oil gavage indicating impaired chylomicron secretion from enterocytes in these mice. This study demonstrated that female *Dennd5b^−/−^* mice are resistant to diet-induced weight gain, hypercholesterolemia, and atherosclerosis ^20^. We also reported that a common single-nucleotide *DENND5B* coding variant, p.(R52K) (rs4930979), was negatively associated with BMI and abdominal circumference in a small human cohort ^20^.

The present study uses *Dennd5b* deficient mice as a model of impaired post-Golgi chylomicron secretion to determine the impact of this condition on body composition and intestinal metabolism and to examine the cellular response to unsecreted chylomicrons. Our findings establish a relationship between *DENND5B* and body fat percent in humans and mice. Furthermore, our data support a key role for *Dennd5b* in chylomicron secretion by enterocytes and demonstrate that post-Golgi chylomicron retention results in activation of autophagy and mitochondrial beta oxidation in the intestinal epithelium. These findings are intriguing because the absorptive intestinal epithelium is not typically considered as highly metabolically active tissue. However, these studies suggest that, under conditions of nutrient accumulation (*e.g*., high-fat diet or impaired chylomicron secretion), enterocytes can activate mitochondrial beta oxidation and this process may be regulated differently in males and females.

## Results

### *DENND5B* is associated with body composition in humans and mice

To examine the impact of DENND5B on body composition in humans, we analyzed data from the Common Metabolic Diseases Knowledge Portal (CMDKP) that compiles genotype-phenotype data from several large genetic datasets ^21-25^. Examination of the impact of individual *DENND5B* SNPs revealed 11 variants significantly associated with BMI. Three variants were positively associated with BMI and 8 were negatively associated suggesting the possibility that both gain and loss-of-function variants exist in the human population (**Fig. 1A**). A gene-level rare variant analysis was performed to examine the impact of predicted loss-of-function mutations on body fat percent. This analysis revealed that rare loss-of-function and missense variants in *DENND5B* are significantly associated with lower body fat percent (**Fig 1B**). To determine if genetic disruption of *Dennd5b* affects metabolic phenotypes in mice, we first measured body weight and body composition of 4-month-old wild type and *Dennd5b^−/−^*mice that had been maintained on a standard chow diet (**Fig. 1C**). *Dennd5b*-deficiency resulted in modestly lower body weight for males, but did not affect body weight of females (**Fig. 1D**). Body composition analysis using echo-MRI revealed that male *Dennd5b^−/−^* mice had significantly lower percent body fat mass compared to wild type mice (**Fig. 1E**). Male *Dennd5b^−/−^* mice had a corresponding increase in percent lean mass compared to wild type controls (**Fig. 1F**). Interestingly, this higher lean mass % was not driven simply by the lower fat mass, but also by a higher absolute lean mass in *Dennd5b^−/−^* males. Body composition data expressed by mass are presented in the Supplementary material (**Supplementary Fig. 1A-C**). As mice aged from 4 to 7 months old, both female and male *Dennd5b^−/−^* mice had slower rates of weight gain compared to their wild type counterparts (**Supplementary Fig. 1D-G**). Percent body weight gain in adult *Dennd5b^−/−^* mice was significantly greater in males compared to females (**Supplementary Fig. 1G**). The differences in body weight gain over 3 months were due to altered body composition, characterized by lower fat mass and higher lean mass in *Dennd5b^−/−^*compared to wild type mice (**Supplementary Fig. 1H,I**).

**Figure 1.**
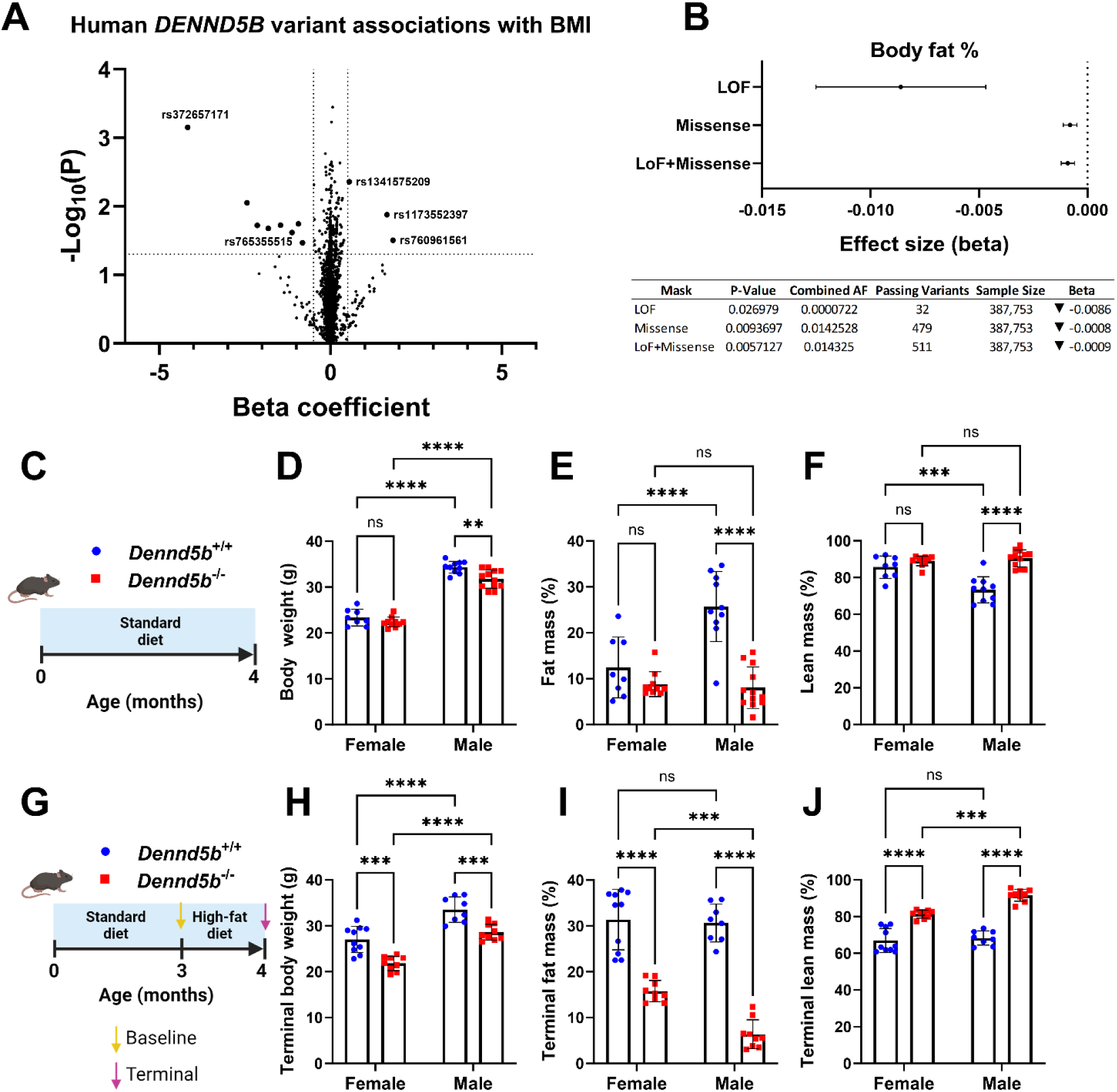
*DENND5B* is associated with body composition in humans and mice. Human gene association data and mouse studies were performed to examine the impact of *DENND5B* on body composition. Data from the Common Metabolic Diseases Knowledge Portal (CMDKP) which compiles genotype-phenotype data from hundreds of large genetic datasets was used to evaluate the association between genetic variation in *DENND5B* and metabolic phenotypes in humans. (A) Volcano plot displaying the effect size and directionality of individual human *DENND5B* single nucleotide polymorphisms (SNPs) effect on BMI. (B) Rare variant analysis for associations of loss-of-function *DENND5B* variants with body fat percent. (C) Schematic of study design to evaluate body composition in 4-month-old *Dennd5b^+/+^* and *Dennd5b^−/−^* female and male mice fed standard chow diet (SD). (D-F) Body weight and echo MRI body composition of 4-month-old SD fed *Dennd5b^+/+^* and *Dennd5b^−/−^* mice. Data are mean +/− standard error (n=8-12/group). Comparisons by Two-way ANOVA with Tukey correction for multiple comparisons. (G) Schematic of study design to evaluate body composition in 4-month-old *Dennd5b^+/+^* and *Dennd5b^−/−^* female and male mice fed high-fat diet. (H-J) Body weights and echo MRI body composition of mice after 1 month of high-fat diet feeding. Data are mean +/− standard error (n=8-10/group). Comparison by Two-way ANOVA with Tukey correction for multiple comparisons. For all statistical comparisons: **p<0.01, ***p<0.001, ****p<0.0001.

To examine the impact of short term high-fat diet (HFD) feeding on body composition, 3-month-old mice were switched from standard rodent diet (18% kcal from fat, 58% carbohydrate, 24% protein) to HFD (45% kcal from fat, 35% carbohydrate, 20% protein) and maintained on this diet for one month (**Fig. 1G**). HFD did not affect expression of *Dennd5b* in intestinal tissue of wild type mice (**Supplementary Fig. 2**). After one month on HFD, both male and female *Dennd5b^−/−^* mice had significantly lower body weight compared to wild type mice (**Fig. 1H**). Male and female wild type mice experienced a body weight increase of about 20%, whereas *Dennd5b^−/−^* mice experienced no weight gain (**Supplementary Fig. 3A-C**). Body composition analysis revealed that both male and female *Dennd5b^−/−^* mice had significantly lower percent fat mass and higher percent lean mass compared to wild type mice (**Fig. 1I,J**). These effects were more prominent in male *Dennd5b^−/−^* mice. No sex differences were observed in the percent fat or lean body mass in wild type mice (**Fig. 1I,J**). Examination of actual body mass changes due to HFD, revealed that lower body weight in *Dennd5b^−/−^* females was due to lower fat mass with similar lean mass. Interestingly, the lower body weights observed in *Dennd5b^−/−^* males were also due to dramatically reduced fat mass, however, this was accompanied by a significant increase in absolute lean mass (**Supplementary Fig. 3D,E).** Together, these findings demonstrate that genetic disruption of *DENND5B* is associated with lower body fat in humans and in mice. In mice, the reduction in body fat is more prominent in males than females. Furthermore, *Dennd5b* disruption in mice increases lean mass, but only in males.

### *Dennd5b*-deficiency prevents postprandial plasma triglyceride elevation after oral oil gavage

Because our previous studies implicate *Dennd5b* in the process of intestinal lipid absorption in mice, we aimed to determine if the differential effect of *Dennd5b^−/−^* on body composition by sex was due to differences in absorption of dietary triglyceride. To address this question, we examined the impact of *Dennd5b*-deficiency on postprandial plasma triglycerides in male and female mice. Consistent with the literature, wild type females had lower postprandial plasma triglyceride concentrations compared to males (**Fig. 2A,B**). Both male and female *Dennd5b^−/−^*mice had a near complete absence of postprandial plasma triglyceride increase. To determine if differences in peripheral lipase activity could be impacting postprandial triglyceride, we measured lipoprotein lipase activity in wild type and *Dennd5b^−/−^* male and female mice before and after injection with heparin to release endothelium-bound lipases. In both genotypes and sexes, post-heparin lipase activity was significantly increased reflecting release of vascular endothelium-bound lipoprotein lipase (**Fig. 2C,D**). Although male *Dennd5b^−/−^* mice had 10% lower post-heparin lipase activity compared to the wild type (**Fig. 2D**), this seems unlikely to account for the reduced post-gavage triglyceride levels. In non-fasting plasma, disruption of *Dennd5b* significantly reduced total cholesterol levels in both sexes, and did not affect plasma triglyceride levels (**Supplementary Fig. 4A-D**). These data indicate that the impact of *Dennd5b* disruption on postprandial plasma triglyceride is similar between sexes and is likely not a result increased peripheral lipolysis.

**Figure 2.**
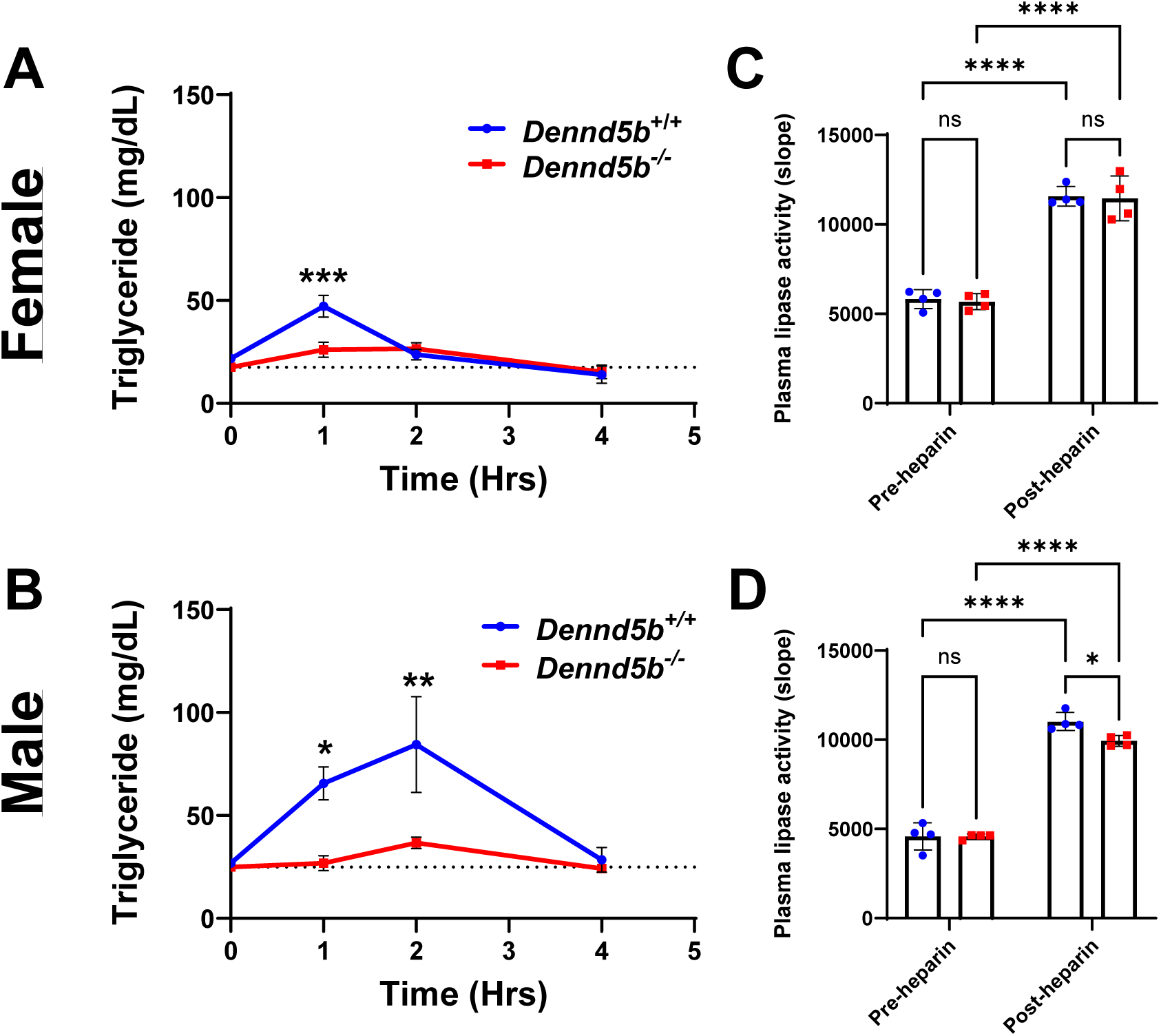
*Dennd5b^−/−^* mice have lower postprandial triglyceride concentrations in plasma. (A-B) *Dennd5b^+/+^* and *Dennd5b^−/−^* mice received an oral gavage of vegetable oil (10 µL/g of body weight) using ball tipped gavage syringe and appearance of plasma triacylglyceride (TAG) was monitored over 4 hours. Both female and male *Dennd5b^−/−^* mice display reduced plasma TAG after gavage compared to *Dennd5b^+/+^* controls. Data are mean +/− standard error (n=4-5/group). Comparison by Two-way ANOVA with Sidak’s multiple comparisons test. (C-D) Plasma lipoprotein lipase activity was measured in plasma that was collected at baseline (pre-heparin) and 10 minutes after retro-orbital injection of heparin (post-heparin, 300 units/kg body weight) in female (C) and male (D) mice. Data are mean +/−standard error (n=4/group). Comparison by Two-way ANOVA with Tukey’s multiple comparisons test. For all statistical comparisons: *p<0.05, **p<0.01, ***p<0.001, ****p<0.0001.

### *Dennd5b*-deficiency results in neutral lipid accumulation along the crypt-villus axis and impaired chylomicron secretion

Because disruption of *Dennd5b* significantly reduced postprandial plasma triglyceride elevations, we next examined the impact of *Dennd5b* on triglyceride absorption in intestinal tissue. Oil red O staining of duodenal sections revealed significant neutral lipid accumulation in the intestinal epithelium of *Dennd5b^−/−^* mice on standard and high-fat diets (**Fig. 3A**). In wild type mice fed standard diet or HFD, small amounts of lipid staining was observed within enterocytes near the villus tip and within the lacteal, reflective of efficient chylomicron secretion (**Fig. 3A,B**). In *Dennd5b^−/−^* intestine, fed standard diet or HFD, large amounts of lipid were observed within enterocytes. Notably, lipid accumulation was observed along the entire length of the villus, but not within the crypts. Intracellular lipid accumulated primarily between the nucleus and the apical surface of the cell, and was less apparent between the nucleus and the basolateral surface (**Fig. 3A,B**). We also observed that small intestine length was significantly increased in *Dennd5b^−/−^* compared to wild type mice (**Supplementary Fig. 5A-D**). Electron microscopy (EM) imaging of duodenal enterocytes six hours after oral oil gavage revealed an accumulation of cytosolic lipid droplets (cLD) and unsecreted chylomicrons within chylomicron secretory vesicles in *Dennd5b^−/−^* tissue (**Fig. 3C**, yellow arrows). Additionally, analysis of mesenteric lymph collected from mice two hours after a duodenal lipid bolus revealed a dramatically reduced concentration of triglyceride in *Dennd5b^−/−^* compared to *Dennd5b^+/+^* lymph (**Fig. 3D**). Taken together, these data demonstrate that *Dennd5b* disruption results in impaired chylomicron secretion and lipid retention in the intestinal epithelium.

**Figure 3.**
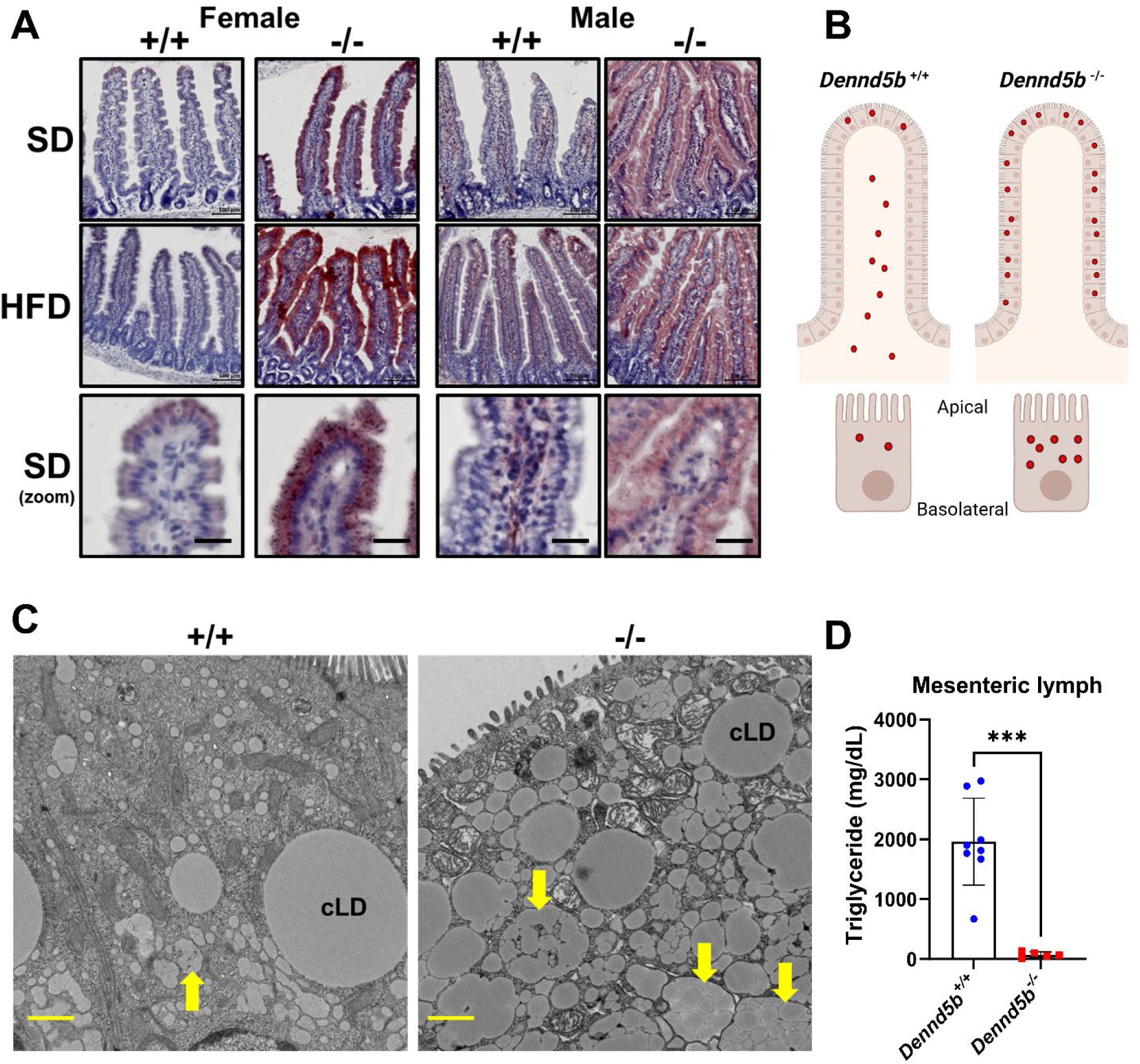
*Dennd5b* deficiency results in neutral lipid accumulation in the small intestine epithelium and impaired chylomicron secretion by enterocytes. (A) Oil red O (ORO) staining of intestinal tissue sections from *Dennd5b^+/+^* and *Dennd5b^−/−^* mice maintained on standard chow diet (SD) or high-fat diet (HFD) for 1 month. (B) Schematic representing differences in the localization of lipid staining in the intestinal epithelium of *Dennd5b^+/+^* and *Dennd5b^−/−^* mice. Red dots indicate neutral lipid localization. (C) Electron microscopy images of duodenal tissue from *Dennd5b^+/+^* and *Dennd5b^−/−^* mice six hours after oral oil gavage. Cytosolic lipid droplets are indicated (cLD) and chylomicron secretory vesicles (yellow arrows). Scale bar = 1 µm. (D) Triglyceride concentration in mesenteric lymph collected from *Dennd5b^+/+^* and *Dennd5b^−/−^* mice two hours after administration of a duodenal bolus of intralipid.

### Disrupted chylomicron secretion increases fecal lipid excretion

To quantitatively assess absorption efficiency of dietary triglyceride, wild type and *Dennd5b^−/−^*mice were fed high-fat diet containing the non-absorbable fatty acid marker sucrose polybehenate. For this study, mice were maintained on standard chow diet until 8 weeks of age then switched to a high-fat diet for 2 weeks. Then, the diet was replaced with and identical diet supplemented with 5% sucrose polybehenate for one week (**Fig. 4A**). At the end of the study, *Dennd5b^−/−^* mice had lower body weight, lower percent fat mass, and higher percent lean mass compared to wild type mice (**Supplementary Fig. 6A-C**). Gas chromatography was used to analyze the ratio of non-absorbable behenic acid to other fatty acids in the feces (and in the diet) and was used to calculate lipid absorption efficiency in these mice. *Dennd5b* disruption significantly reduced fatty acid absorption efficiency in both sexes (**Fig. 4B**). This effect was more prominent in males than in females (males −7.9% and females −3.7% compared to their respective wild type controls, p<0.01). The consequent impact on fecal fatty acid excretion in *Dennd5b^−/−^* mice was a 2.6-fold increase in females (p<0.01) and a 4-fold increase in males (p<0.0001) (**Fig. 4C**). Analysis of the relationship between fatty acid absorption efficiency and body fat mass showed a positive linear relationship for males, but not for females, (**Fig. 4D,E**) demonstrating that this relationship is impacted by biological sex. The mRNA levels of key enteroendocrine hormones involved in fat digestion, cholecystokinin (*Cck*) and secretin (*Sct*), were not altered in *Dennd5b^−/−^* mouse intestinal tissue suggesting that these are not a major contributor to the effects on fatty acid absorption (**Supplementary Fig. 7A,B**). Overall, these findings demonstrate that disruption of chylomicron secretion has a more prominent impact on fatty acid absorption efficiency in males than in females and that males have a stronger correlation between dietary fatty acid absorption efficiency and body fat mass.

**Figure 4.**
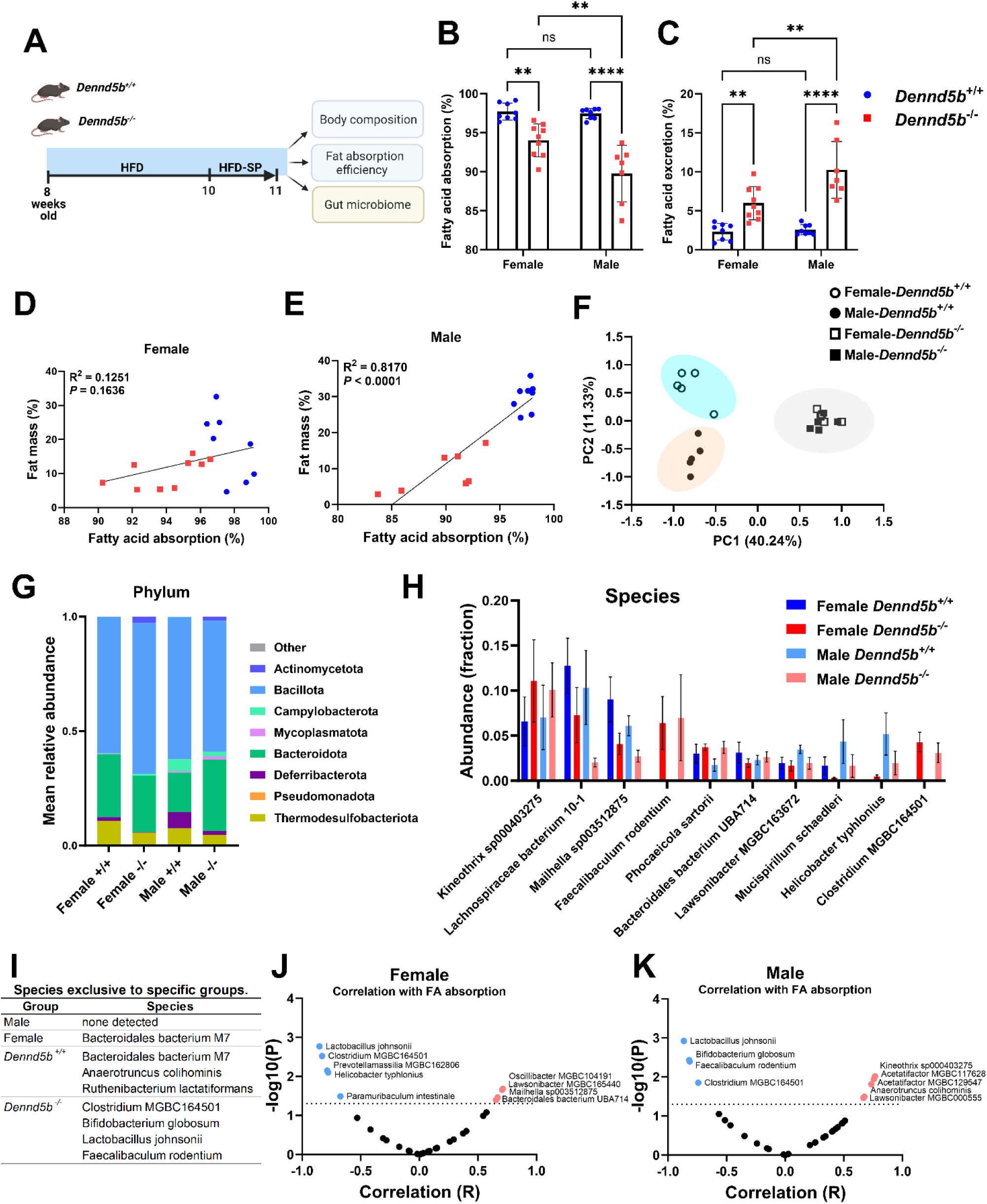
*Dennd5b^−/−^* mice have reduced absorption of dietary triglycerides and altered gut microbiome. (A) Schematic representing the study design. *Dennd5b^+/+^* and *Dennd5b^−/−^* mice of both sexes were maintained on standard chow diet until 8 weeks of age and then switched to high-fat diet (HFD) for 2 weeks. Following two weeks acclimation, HFD was replaced with an identical diet supplemented with 5% sucrose polybehenate (HFD-SP). (B-C) Dietary fatty acid absorption efficiency and Fecal fatty acid excretion. Data are mean +/− standard error (n=7-9/group). Comparison by Two-way ANOVA with Tukey’s multiple comparisons test. (D-E) Linear regression analysis of the relationship between fatty acid absorption and body fat % revealed a significant relationship in males (E), but not in females (D). (F) Principal components analysis shows significant global changes in gut microbiome. *Dennd5b^+/+^* females and males cluster separately to the left, while *Dennd5b^−/−^* females and males cluster together to the right. n=5/group for gut microbiome studies. (G) Differential abundance of microbiota at phylum level in *Dennd5b^+/+^* and *Dennd5b^−/−^*female and male fecal samples. (H) Comparison of the relative abundance of the top 10 most abundant gut microbiota species. (I) Microbiome species detected in the fecal samples exclusively in one sex or genotype. (J, K) Correlation analysis of the relationship between abundance of individual microbiome species and fatty acid absorption in females (J), and in males (K). For all statistical comparisons: **p<0.01, ****p<0.0001.

### Disrupted chylomicron secretion alters the gut microbiome

Our findings of increased fecal fatty acid excretion suggest that fatty acid transit through the distal intestine is increased. Based on this, we hypothesized that the elevated fatty acid content would impact the gut microbiome. Fresh fecal samples were collected from HFD fed wild type and *Dennd5b^−/−^* mice and microbiome analysis was performed. Principal components analysis of fecal microbe abundance revealed a strong separation between wildtype and *Dennd5b^−/−^* mice, as well as separation between male and female wildtype mice that was lost in *Dennd5b^−/−^* animals (**Fig. 4F**). At the Phylum level, several microbiome shifts in *Dennd5b^−/−^* mice were evident including expansion of Actinomycetota and contraction of Deferribacterota and Thermodesulfobacteriota populations (**Fig. 4G**). Campylobacterota appeared to be more abundant in males than females regardless of genotype. At the species level, comparison of the top 10 most abundant microbes demonstrated that *Dennd5b*-deficiency revealed striking changes to the gut microbiome (**Fig. 4H**). This includes the appearance of two microbe species that are not detected in wild type mice, Faecalibaculum rodentium and Clostridium MGBC164501, and the expansion of Kineothrix. All of these effects were consistent in both male and female *Dennd5b^−/−^* mice. Other species, including Lachnospiraceae, Mailhella, and Mucispirillum schcaedleri, appear to be less abundant in *Dennd5b^−/−^* mice. Species detected exclusively in one sex or genotype are listed in **Fig. 4I**. To determine if altered gut microbiome contributes to differences in fatty acid absorption, we performed a correlation analysis to examine the relationship between abundance of individual microbe species and fatty acid absorption efficiency in males and females. Several species were negatively correlated with fatty acid absorption including two that were consistent between male and female mice: Lactobacillus johnsonii and Clostridium MGBC164501 (**Fig. 4J,K**). Several species were positively correlated with fatty acid absorption including Kineothrix, Oscillibacter, and Lawsonibacter; however, Lawsonibacter was the only species consistently associated in both sexes. In summary, genetic disruption of *Dennd5b* in mice fed HFD results in a striking expansion of specific microbe species and a loss of features that distinguish wild type mice by biological sex. Correlations for some species suggest possible roles in modulating dietary fatty acid absorption.

### Gene expression profiles in *Dennd5b^−/−^* intestine indicate altered fatty acid metabolism

To examine the global impact of intracellular lipid accumulation in *Dennd5b^−/−^* enterocytes, we performed RNA sequencing on duodenum tissue from mice maintained on standard chow or high-fat diet. Principal components analysis revealed that *Dennd5b* deficiency amplifies differences in intestinal gene expression in response to high fat diet compared to wild type mice (**Fig. 5A**). Volcano plots were used to visualize differentially expressed genes (DEGs) in chow-fed mice (**Fig. 5B**). Male mice had a greater number of total DEGs in response to *Dennd5b* disruption compared to female mice (2,486 vs 785 DEGs, respectively). DEGs were roughly evenly distributed between upregulation and downregulation in both sexes. Gene Ontology Biological Process analysis of DEGs revealed significant upregulation of genes involved in fatty acid metabolism and mitochondrial function (**Fig. 5C**). (**Supplementary Fig. 8A-D**)

**Figure 5.**
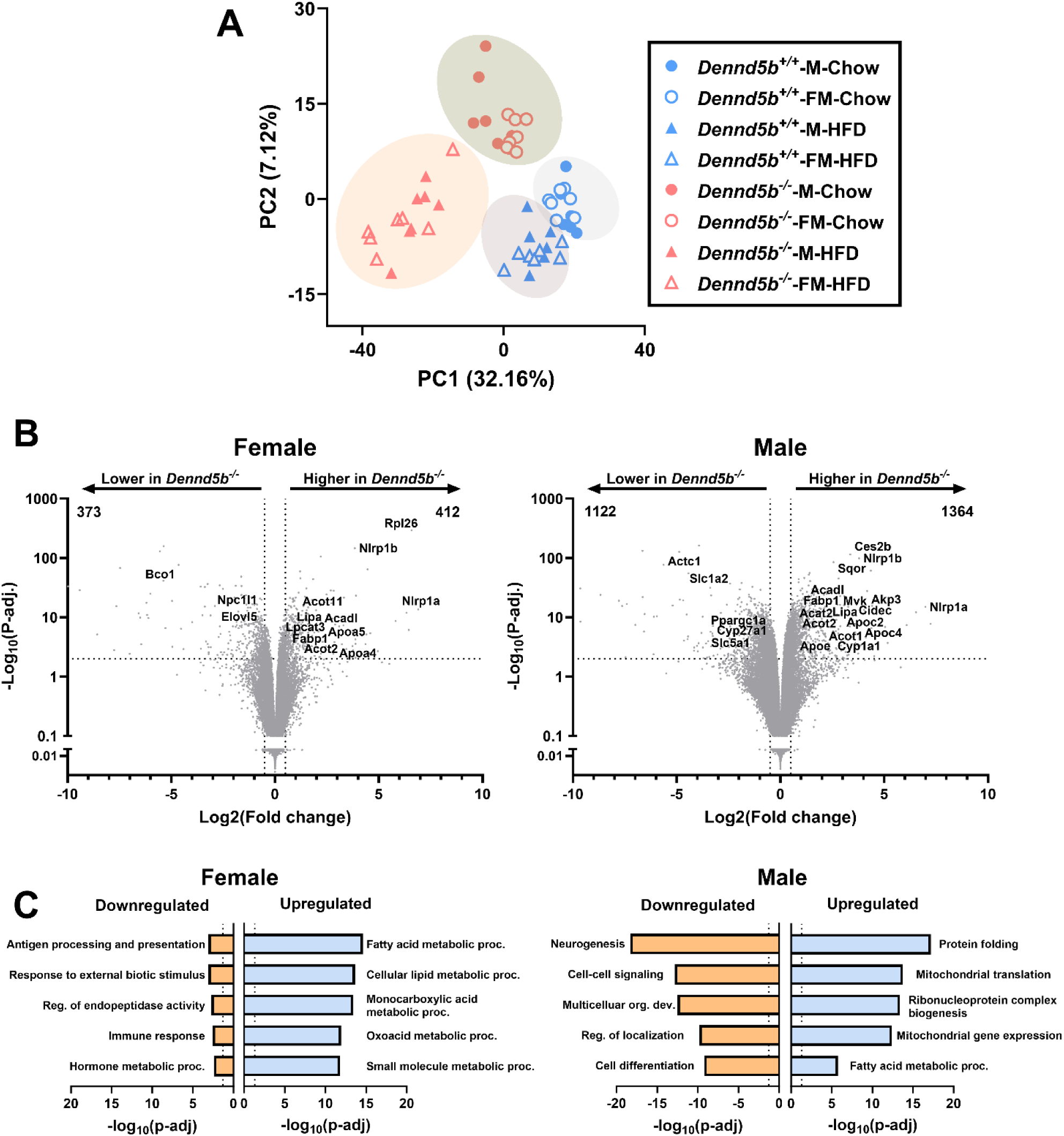
Gene expression profiles in *Dennd5b^−/−^* intestine indicate altered fatty acid metabolism. Bulk RNAseq analysis was performed on duodenum tissue from wild type and *Dennd5b^−/−^*mice fed standard chow or high-fat diets. (A) Principal components analysis. (B) Volcano plots displaying differentially expressed genes as a result of *Dennd5b* disruption in male and female mice fed standard chow diet. (C) Gene ontology Biological Process pathway enrichment analysis on differentially expressed genes to identify up and downregulated pathways in response to *Dennd5b* disruption in male and female mice fed standard chow diet.

### Mitochondrial beta oxidation is increased in intestinal tissue on high-fat diet and when chylomicron secretion is impaired

To understand how fatty acids are being metabolized by intestinal tissue in the context of impaired chylomicron secretion, we compared metabolic function of intestinal tissue from wild type and *Dennd5b^−/−^* mice. First, we examined intestinal mRNA abundance of genes involved in fatty acid metabolism. In wild type mice, HFD increased *Ppara*, strongly inducing expression of genes involved in beta oxidation and ketogenesis (**Fig. 6A**). In *Dennd5b^−/−^* mice, *Ppara* is less strongly induced and even downregulated in males. Despite this, beta oxidation genes are increased in *Dennd5b^−/−^* compared to wild type on both SD and HFD (**Fig. 6A and Supplementary Fig. 9**). Notably, markers of fatty acid uptake and ketogenesis are strongly repressed in *Dennd5b^−/−^* intestine. This may be mediated by concurrent activation of *Pparg* in the knockouts and also by increased mitochondrial and endoplasmic reticulum stress (**Fig. 6A,B**). Electron microscopy imaging of duodenal epithelium revealed altered mitochondrial morphology in *Dennd5b^−/−^* enterocytes, consistent with increased mitochondrial activity and mitochondrial stress (**Fig. 6C**).

**Figure 6.**
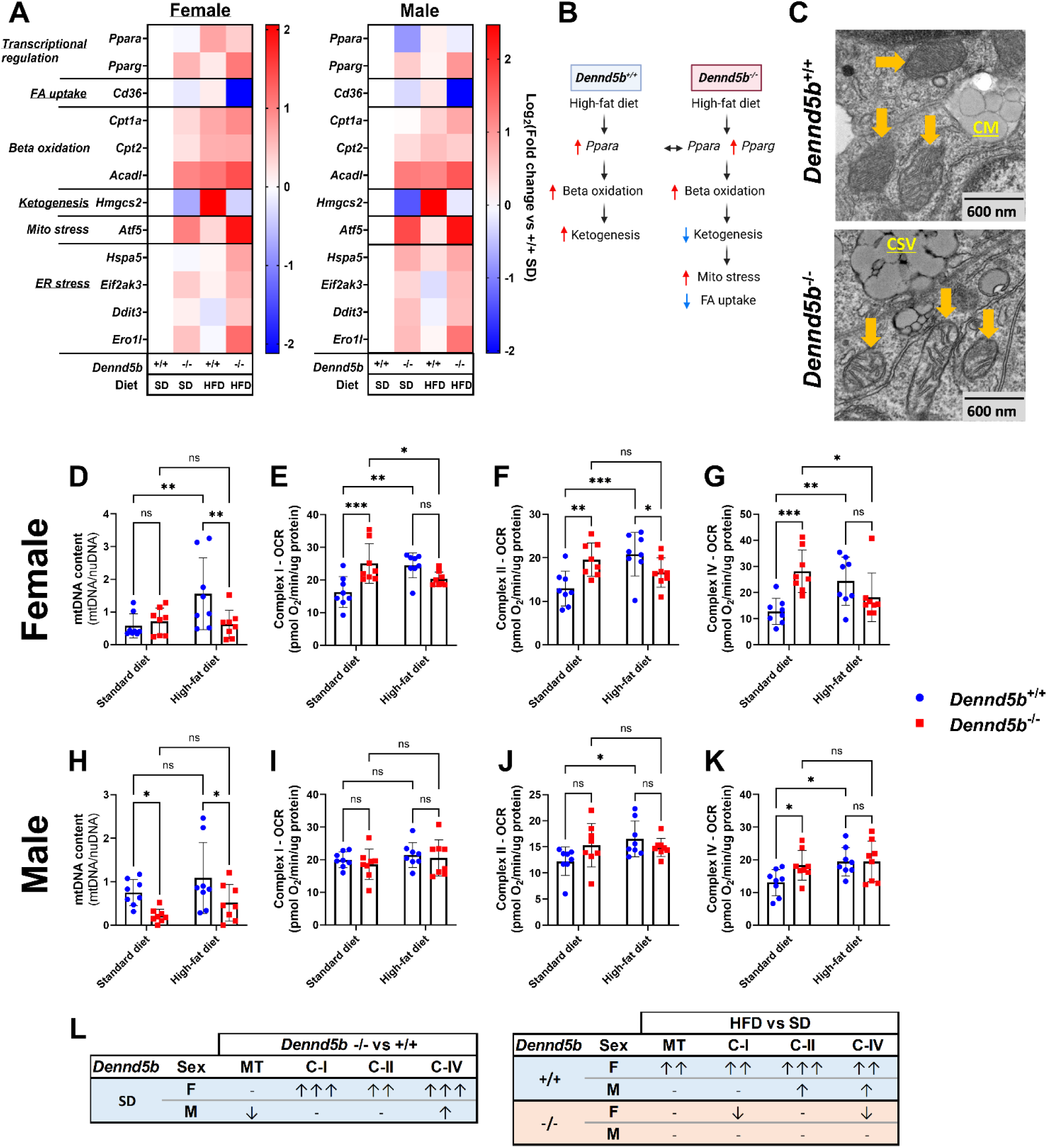
Fatty acid oxidation is induced in intestinal tissue by high-fat diet and impaired chylomicron secretion. (A) Duodenal mRNA abundance of genes involved in the beta oxidation pathway revealed increased expression of these genes in *Dennd5b^−/−^* mice compared to wild types in both females and males. (B) Summary of effects of *Dennd5b^−/−^*on fatty acid metabolic pathway. (C) Transmission electron micrographs of mouse duodenum. Yellow arrows indicate mitochondria. CM–chylomicron (secreted to intercellular space), CSV–chylomicron secretory vesicles. (D, H) Mitochondrial density in the intestinal tissues from 4-month-old female and male *Dennd5b^+/+^* and *Dennd5b^−/−^*mice maintained on standard chow diet (SD) or high-fat diet (HFD) for 1 month. Data are mean +/− standard deviation (n=8/group). Comparison by Two-way ANOVA with uncorrected Fischer’s LSD multiple comparisons test. (E-G, I-K) Respirometry measurements of complex I, II, and IV in females (E-G) and males (I-K) *Dennd5b^+/+^* and *Dennd5b^−/−^* mice maintained on SD or HFD for 1 month. Data are mean +/−standard deviation (n=8/group). Comparison by Two-way ANOVA with uncorrected Fischer’s LSD multiple comparisons test. (L) Summary table of mitochondrial responses to HFD feeding from panels D-K. For all statistical comparisons: *p<0.05, **p<0.01, ***p<0.001.

We next examined mitochondrial content and function in mouse intestinal tissue. In wild type mice, HFD feeding increased mitochondrial density in females, but not males (**Fig. 6D,H**). *Dennd5b* deficiency resulted in reduced mitochondrial DNA content in male intestine on both SD and HFD. However, *Dennd5b* disruption only affected mitochondrial density in females when fed HFD, as the lack of *Dennd5b* appeared to prevent HFD-induced increase in mtDNA (**Fig. 6D,H**). The lower mitochondrial content in *Dennd5b^−/−^* may be mediated in part by reduced expression of the mitochondrial biogenesis regulating transcription factor *Ppargc1a* (**Supplementary Fig. 10**). We assessed mitochondrial function by performing respirometry analysis on intestinal tissue. In wild type mice, HFD feeding increased intestinal mitochondrial respiration and this effect was more prominent in females than males (**Fig. 6E-G, I-K**). *Dennd5b* deficiency significantly increased intestinal mitochondrial activity in SD fed mice. This effect was also more prominent in females and was not further impacted by HFD feeding. Interestingly, despite a dramatic drop in intestinal mitochondrial density of male *Dennd5b^−/−^*mice, these mice maintained similar complex IV activity compared to wild type males (**Fig. 6I-K**). The impact of *Dennd5b* and diet on mitochondrial content and function are summarized in **Fig. 6L**. Normalizing mitochondrial function measurements to mitochondrial DNA revealed that *Dennd5b^−/−^* intestine had higher respiratory activity per mitochondrion compared to wild type, particularly on SD (**Supplementary Fig. 11**). To investigate the contribution of lipid droplet-derived fatty acids to mitochondrial substrates in *Dennd5b^−/−^* intestine, we examined the expression of the lipid droplet coat protein (*Plin2*) and lipases involved in lipid droplet hydrolysis (*Pnpla2* and *Lipe*). *Plin2* expression increased in response *Dennd5b* disruption, while both lipases were significantly downregulated (**Supplementary Fig. 12**). Collectively these data demonstrate increased mitochondrial beta oxidation of fatty acids and suggest that the source of FAs is not from triglyceride stored in lipid droplets, raising the possibility that unsecreted chylomicrons may be a source of FAs.

### Autophagy is increased in *Dennd5b^−/−^* intestine tissue

Electron micrographs of fasting intestinal tissue revealed the presence of large, electron-dense structures within enterocytes of *Dennd5b^−/−^* mice (**Fig. 7A**). These membrane-bound structures contained cytoplasm and membranous whorls characteristic of autophagic degradation of membrane-bound structures. These were not observed in the intestinal epithelium of wild type mice. We speculated that these may be autophagosomes and hypothesized that autophagy may be a mechanism by which *Dennd5b^−/−^* enterocytes dispose of accumulated chylomicron secretory vesicles and damaged mitochondria. Western blotting of intestinal lysates for the autophagy marker LC3 showed a significant increase in total LC3-II protein as well as an increase in the LC3-II to LC3-I ratio indicating increased activation of autophagy in *Dennd5b^−/−^*mice compared to wild type (**Fig. 7B**). P62 protein levels were not different between wild type and *Dennd5b^−/−^* mice, however, we also observed that mRNA levels of *Sqstm1* (gene encoding P62) were elevated in *Dennd5b^−/−^* mice, suggesting a possible upregulation of this protein that may be offsetting its degradation by autophagy (**Fig. 7B and Supplementary Fig. 13A**). Additionally, mRNA abundance of key enzymes involved in the initiation of autophagy, *Atg5* and *Atg16l1*, increased in *Dennd5b^−/−^* intestinal tissue (**Supplementary Fig. 13B,C**). Immunofluorescence staining of duodenal tissue revealed the presence of cytosolic LC3-positive autophagosomes only in *Dennd5b^−/−^* mice (**Fig. 7D**). When *Dennd5b^−/−^* mice were switched to a diet that contains no fat, LC3-positive autophagosomes were no longer detected in intestinal tissue indicating that autophagy in these mice is induced by dietary lipid. Lipophagy is a specialized form of autophagy characterized by the liberation of free fatty acids from triglyceride for utilization in beta oxidation. In this process, triglyceride is hydrolyzed by lysosomal acid lipase (*Lipa*). We found that *Lipa* expression is significantly increased in *Dennd5b^−/−^* mice (**Fig. 7E**) and that expression of *Lipa* is strongly correlated with the autophagy activating enzyme *Atg5* (**Fig. 7F**).

**Figure 7.**
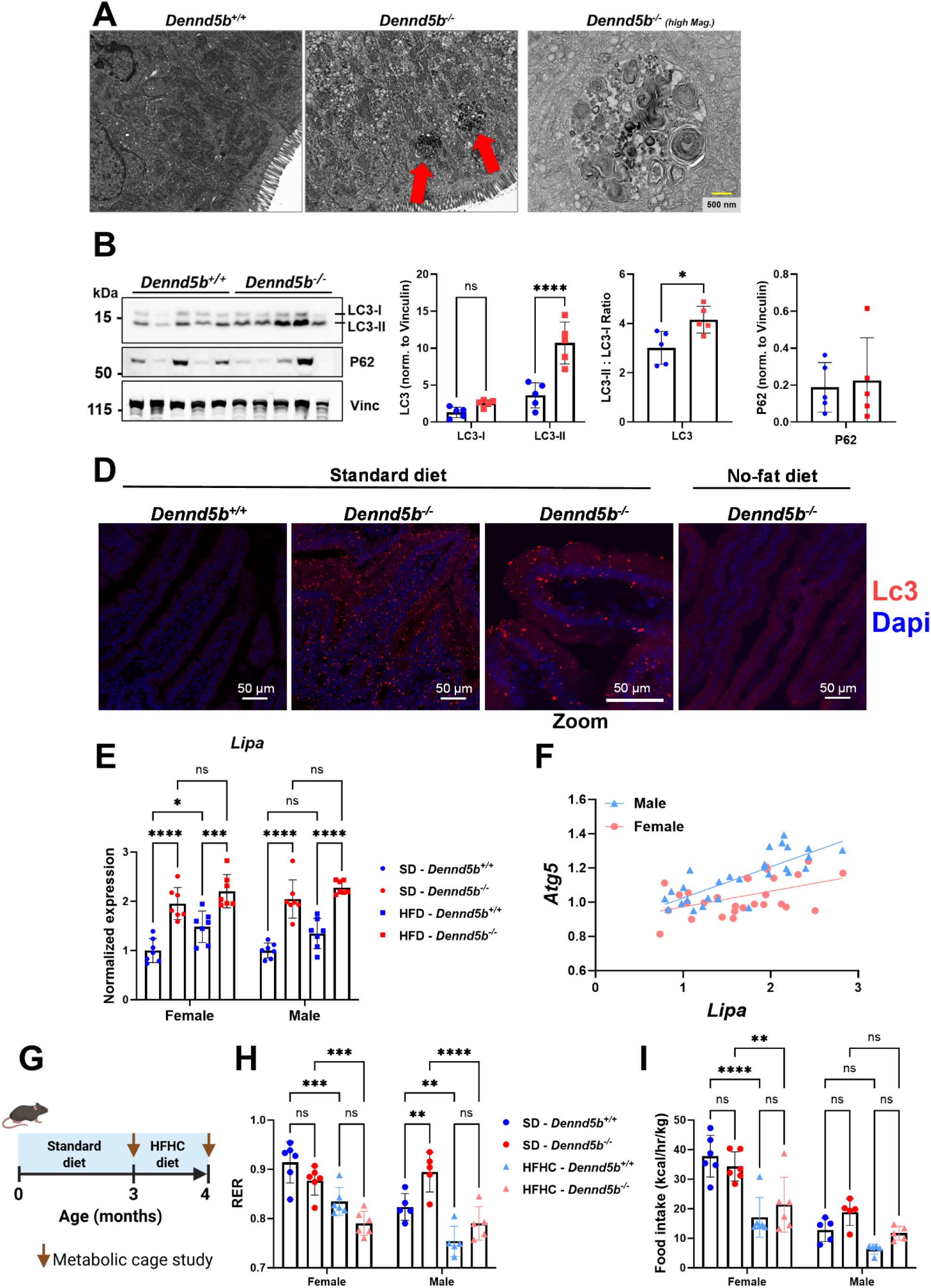
Autophagy is increased in *Dennd5b^−/−^* intestine. (A) TEM images of *Dennd5b^+/+^* and *Dennd5b^−/−^* duodenal enterocytes previously published by Gordon et al, are unedited and available for reuse under Creative Commons Attribution 4.0 International License https://creativecommons.org/licenses/by/4.0/. Large electron dense intracellular structures were observed in *Dennd5b^−/−^* (middle), but not wildtype (left) enterocytes. Higher magnification imaging of these structures (right) reveals a membrane border and complex inner structure with membrane “whorls” characteristic of autophagosomes. (B) Western blots of intestinal tissue from *Dennd5b^+/+^* and *Dennd5b^−/−^* mice on standard chow diet. (C) Normalized duodenal LC3-II protein abundance. Data are mean +/− standard error (n=4/group). Comparison by Two-way ANOVA with Tukey’s multiple comparisons test. (D) Immunofluorescence staining for Lc3 in intestinal tissue from *Dennd5b^+/+^* and *Dennd5b^−/−^* mice on SD or no-fat diet. (E) Relative abundance of *Lipa* mRNA in female and male *Dennd5b^+/+^* and *Dennd5b^−/−^* mice maintained on SD or HFD for 1 month. Data are mean +/− standard error (n=7/group). Comparison by Two-way ANOVA with Tukey’s multiple comparisons test. (F) Linear regression analysis of the relationship between the mRNA abundance of *Atg5* and *Lipa* revealed a positive correlation in both males and females. (G) Wildtype and *Dennd5b^−/−^* mice underwent metabolic testing by indirect calorimetry (CLAMS) at baseline on standard diet (circles) and the same mice were remeasured after 1 month on high-fat high-cholesterol (HFHC) diet (42% kCal fat + 0.2% cholesterol) (triangles). (H) *Dennd5b^−/−^* mice exhibit a downward shift in respiratory exchange ratio (RER) in response to HFHC diet similar to *Dennd5b^+/+^* mice. (I) Food intake of mice during the metabolic cage study. Comparison by two-way ANOVA with Tukey post-hoc test. For all statistical comparisons: *p<0.05, **p<0.01, ***p<0.001, ****p<0.0001.

To examine fatty acid oxidation in the context of the whole animal, we performed metabolic cage studies on wild type and *Dennd5b^−/−^* mice. These data reveal that even though *Dennd5b^−/−^* mice do not absorb dietary fatty acids, they experience a significant reduction in respiratory exchange ratio (RER) when switched from standard diet to a high-fat diet (**Fig. 7G,H**). No genotype mediated differences in food intake were observed (**Fig. 7I**). Taken together, these findings support the hypothesis that lipophagy is upregulated in the intestinal epithelium in the context of impaired chylomicron secretion and that this may contribute to increased fatty acid oxidation and mitochondrial stress in this context (**Fig. 8**).

**Figure 8.**
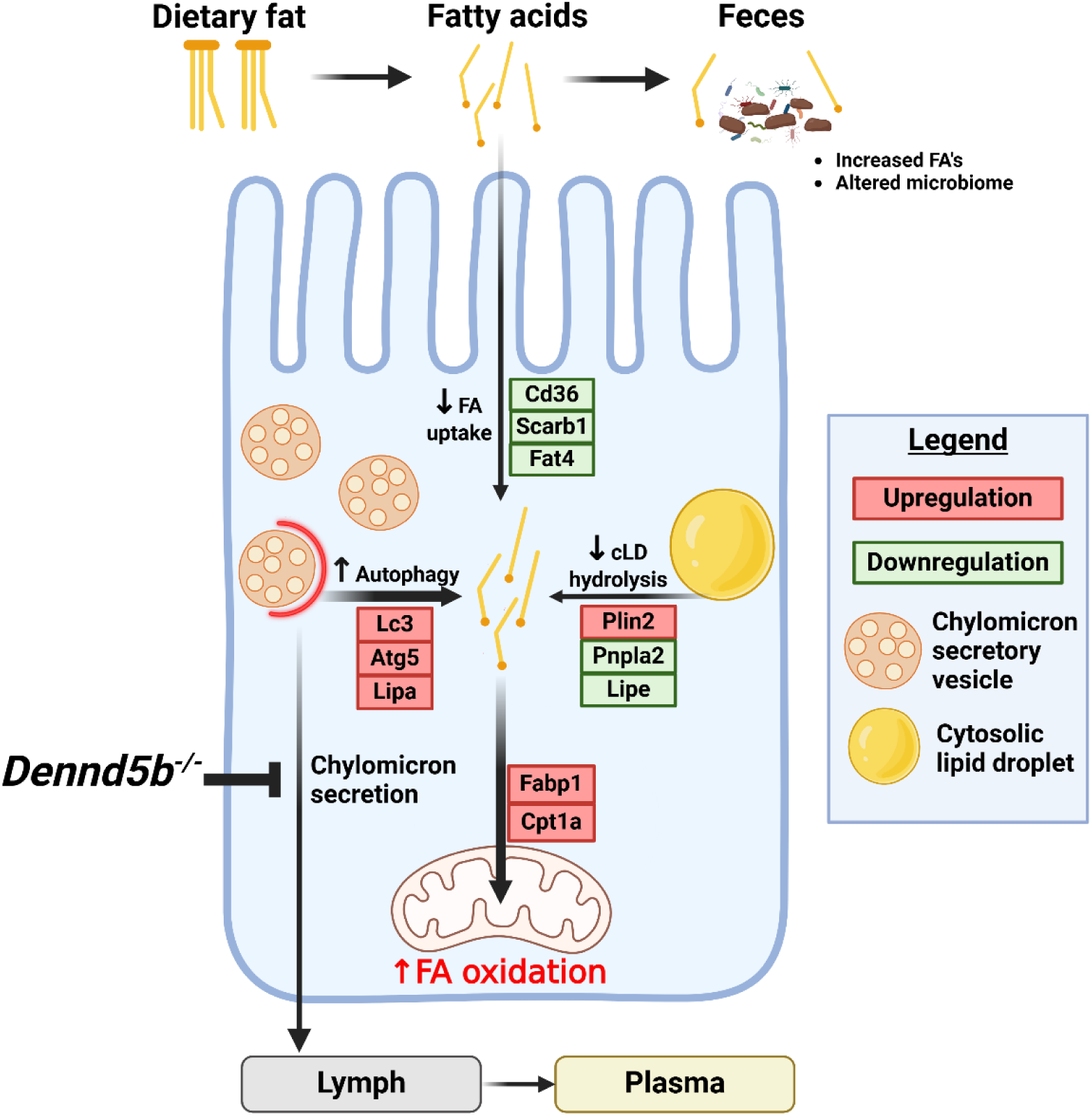
Summary of cellular metabolic response to impaired chylomicron secretion by genetic disruption of *Dennd5b*. Genetic disruption of *Dennd5b* in mice results in impaired chylomicron secretion and accumulation of unsecreted chylomicrons in the intestinal epithelium. Our findings demonstrate a cellular response to unsecreted chylomicron accumulation characterized by 1) downregulation of fatty acid (FA) uptake and increased FA excretion, 2) downregulation of cytosolic lipid droplet (cLD) hydrolysis, 3) upregulation of autophagy, and 4) increased FA oxidation.

## Discussion

Our findings reveal that genetic disruption of *DENND5B* results in altered body composition characterized by reduced body fat in humans and in mice. In mice, reduced fat mass was observed in both sexes and increased lean mass only in male mice. These metabolic phenotypes appear to be driven by impaired intestinal absorption of dietary lipid due to defective secretion of formed chylomicrons by enterocytes. This results in an accumulation of neutral lipid within the intestinal epithelium as both lipid droplets and unsecreted chylomicrons. Saturation of the intestinal epithelium with lipid and downregulation of fatty acid importers contribute to increased excretion of fecal fatty acids and an altered gut microbiome. Ultimately, the cellular accumulation of unsecreted triglyceride in the *Dennd5b^−/−^* epithelium results in induction of autophagy and mitochondrial oxidation of accumulated fatty acids.

Our data demonstrate sex-biased differences in the impact of *Dennd5b* on body weight and body composition. In mice, the magnitude of these effects was often greater in males compared to females. Disruption of intestinal chylomicron secretion in *Dennd5b^−/−^* mice, despite preventing the appearance of ingested triglyceride in the plasma of both sexes, more prominently affected body weight and percent body fat in males. An effect on females was only observed when fed HFD. An important discovery related to the changes in body composition of *Dennd5b^−/−^* mice is that, specifically in male mice, there was an increase in absolute lean mass compared to wild type mice. This suggests that, in addition to limiting uptake of dietary triglyceride, additional mechanisms may be influenced in this context which promote accumulation of lean mass. This is particularly interesting because there are very few known genetic situations which effectively increase lean mass. When comparing the differences between the impact of *DENND5B* on metabolic phenotypes in humans and mice, it is important to consider that the probability of loss-of-function intolerance for human *DENND5B* mutations is very high, according to the gnomAD database (version 4.0.0) ^26^. This suggests that complete loss-of-function mutations are not well tolerated or may be incompatible with life in humans. Thus, the human SNP effects are likely driven by point mutations with more modest effects on protein expression or function. This may explain the relatively modest effects in humans compared to the prominent metabolic phenotypes observed in the *Dennd5b^−/−^* mouse model utilized in the presented studies.

Our findings suggest that the observed metabolic phenotypes in mice are likely driven by impaired intestinal lipid absorption in the *Dennd5b^−/−^* mice. Our postprandial plasma triglyceride studies are consistent with the literature demonstrating that wild type males have higher postprandial triglyceride levels compared to females ^27,28^. For both sexes, disruption of *Dennd5b* results in significantly blunted increases in postprandial plasma triglyceride. Measurements of pre and post-heparin plasma lipase activity suggest that neither the sex or genotype-based differences in postprandial plasma triglyceride are due to differences in peripheral lipase activity. Histology examination of intestinal tissue demonstrated significant neutral lipid accumulation in the intestinal epithelium and EM imaging of enterocytes revealed an abundance of formed chylomicrons in secretory vesicles persisting even 6 hours after an oral oil gavage. A detailed time series of EM images of *Dennd5b^−/−^* enterocytes after oil gavage can be found in our previous publication ^20^. Additionally, a dramatic reduction in triglyceride concentrations was observed in intestinal lymph after a lipid bolus in *Dennd5b^−/−^* mice compared to wild type. All of these findings support the conclusion that genetic disruption in *Dennd5b* in mice results in defective chylomicron secretion by intestinal epithelial cells. Studies with the non-absorbable fatty acid marker sucrose polybehenate revealed that wild type females and males have similar efficiency of dietary fatty acid absorption over time. In both sexes, *Dennd5b^−/−^* mice experienced a drop in fatty acid absorption efficiency and corresponding increase in fecal fatty acid excretion, although this effect was greater in males compared to females. Here, we report a strong positive association between dietary fatty acid absorption efficiency and body fat mass, however, this association was only present in male mice. While it seems intuitive that more fat absorbed ultimately leads to more fat stored, the lack of association in females suggests a possible disconnect in this process.

Gene expression profiles in absorptive epithelial cells change as you move from the crypt toward the villus tip resulting in preferential absorption of specific nutrients within specific regions ^29^. Under normal conditions, lipids are preferentially absorbed by enterocytes near the villus tip. This is apparent in our Oil red O staining of small intestine tissue from wild type mice fed SD or HFD. As expected, neutral lipid staining was observed within the lacteals of wild type mice. This is indicative of successful chylomicron secretion. Genetic disruption of *Dennd5b* resulted in striking accumulation of neutral lipid within intestinal epithelial cells spanning from just above the crypt all the way up to the villus tip. This uncharacteristic accumulation of lipid within enterocytes very close to the base of the villus suggests possible reprogramming of absorptive enterocytes along the crypt villus axis. One possible explanation for this effect is that it could be an adaptive mechanism to increase bulk fatty acid uptake under the condition of inefficient dietary lipid absorption. The small intestine also has the ability to shorten or elongate in response to the nutritional status of the animal. Our observation of elongation of the small intestine in *Dennd5b^−/−^* mice may be a complementary or parallel mechanism with enterocyte reprogramming in an attempt to increase the total mass of lipid absorbed by the animal when this process is inefficient or impaired. We also observed that accumulated lipid within *Dennd5b^−/−^* enterocytes had a distinctive subcellular localization, concentrating specifically in the space between the apical membrane and the nucleus. This localization is consistent with the positioning of the Golgi, while the lack of detectable lipid near the basolateral membrane is consistent with the hypothesis that Dennd5b protein plays a role in Golgi to plasma membrane trafficking of chylomicron secretory vesicles possibly by acting as a guanine nucleotide exchange factor for GTPase proteins involved in vesicular trafficking ^20,30,31^.

Several cell types and tissues have been demonstrated to upregulate autophagy to deal with accumulation of excess or unneeded molecules. Lipophagy is a specific form of autophagy where the target for degradation is usually lipid droplets or lipid loaded vesicles. During lipophagy, intracellular neutral lipids are hydrolyzed by lysosomal acid lipase (Lipa) to generate free fatty acids which can be utilized in mitochondrial beta oxidation. We found that disruption of chylomicron secretion in *Dennd5b^−/−^* mice resulted in increases in markers of autophagy in the intestinal tissue, both at the protein and RNA level. Visual confirmation of autophagy was provided by the presence of LC3-positive autophagosomes detected by immunofluorescence in *Dennd5b^−/−^* intestinal tissues. The absence of LC3-positive staining in *Dennd5b^−/−^* mice fed a no-fat diet supports our hypothesis that dietary lipid is activating autophagy in these mice. The increase in autophagy markers also coincided with increased expression of *Lipa*, supportive of our conclusion that *Dennd5b^−/−^* enterocytes activate lipophagy as a mechanism to dispose of unsecreted chylomicrons and prevent cytotoxicity. Furthermore, the fact that *Dennd5b^−/−^* mice experience a downward shift in RER in response to high-fat diet feeding indicates that despite the inability to absorb dietary fatty acids beyond the intestinal epithelium, the mice are still utilizing some dietary fatty acids.

Genetic disruption of genes involved in intestinal lipid absorption has been previously reported to impact intestinal tissue triglyceride content and beta oxidation. One study from Iqbal et al. demonstrated in mice that genetic disruption of *Mttp*, a protein responsible for the initial lipidation of Apob, resulted in lipid accumulation in the intestinal epithelium, increased expression of genes involved in fatty acid oxidation, and increased beta oxidation in intestinal tissue ^32^. Another study, from Auclair et al., demonstrated that mice with heterozygous disruption of *Sar1b*, a protein responsible for ER to Golgi trafficking of chylomicrons, have increased levels of proteins involved in beta oxidation in intestinal tissue ^33^. Our studies expand on these findings by examining a distinct downstream step of the chylomicron secretion pathway, directly measuring tissue mitochondrial function, establishing the mechanistic link with lipophagy, and by revealing the importance of biological sex in the metabolic response. We show that HFD increases mitochondrial density in wild type female but not male intestine. Wild type females also experienced a robust increase in intestinal mitochondrial activity in all three complexes measured while this effect was much less pronounced in males, with more modest increases and in only 2 of the complexes measured. When chylomicron secretion was impaired by disruption of *Dennd5b,* standard diet fed mice increased intestinal mitochondrial function compared to wild type controls. Again, this effect was of a greater magnitude in females compared to males. *Dennd5b^−/−^* prevented HFD-induced mitochondrial expansion in females and actually reduced mitochondrial content in males regardless of diet. It is important to consider that all of the mitochondrial function measurements are normalized by total protein. Therefore, if we account for the reduced mitochondrial content in *Dennd5b^−/−^* males, they actually have much higher mitochondrial activity on a per mitochondria basis. Upregulation of ketogenesis is a normal physiological response to high levels of beta oxidation, while this effect was observed in wild type mice, *Hmgcs2* expression was significantly downregulated in *Dennd5b^−/−^* mice coinciding with increased markers of mitochondrial stress. Taken together, these data demonstrate a sex difference in the normal metabolic response to HFD with female intestinal tissue more robustly increasing mitochondrial activity in response to diet. This biological sex effect is lost when lipid absorption is disrupted in *Dennd5b^−/−^* mice. It is not yet clear to what extent this pathway may be induced in individuals with genetic variants of *DENND5B* or other genes involved in chylomicron secretion.

In conclusion, these studies add to our mechanistic understanding of the role that *Dennd5b* plays during intestinal absorption of dietary lipids. Our findings reveal novel sex-biased differences in the impact of disrupted chylomicron secretion on body phenotypes and intestinal metabolic function. Furthermore, our studies uncovered a mechanism by which enterocytes can break down unsecreted chylomicrons through lipophagy and utilize their fatty acids in mitochondrial beta oxidation. Thus, these studies emphasize the potential contribution of biological sex-biased differences in the intestinal lipid absorption pathway to metabolic phenotypes. Future studies on genetic and/or environmental influences on this process may contribute to a mechanistic explanation for postprandial lipid and body composition differences among individuals with similar lifestyles. A better understanding of how this process is regulated could provide information that is useful for developing novel strategies to prevent obesity and atherosclerotic cardiovascular disease.

## Data availability statement

The data that support the findings of this study are available on request from the corresponding author.

## Acknowledgements and grant support

This work was supported by National Institutes of Health National Institute of Digestive, Diabetes and Kidney Diseases R01DK133184 (S.M.G) and the National Institute of General Medical Sciences P30GM127211 (Pilot award to S.M.G). The content is solely the responsibility of the authors and does not necessarily represent the official views of the National Institutes of Health. This work was also supported by an American Heart Association Predoctoral Fellowship to O.K.H (24PRE1195834).

We would like to acknowledge the Light Microscopy Core Facility at the University of Kentucky College of Medicine for training lab personnel and for access to imaging equipment.

Special acknowledgement to Dr. Edward Neufeld who acquired electron microscopy images.

## Materials and Methods

### Mouse housing

Mice were housed at the University of Kentucky Division of Laboratory Animals Resources in individually ventilated cages, with a maximum of 5 mice per cage. The mice were housed with a light: dark cycle of 14 hours of light and 10 hours of darkness, and the ambient temperature was maintained at 22°C (72°F). Teklad Sani-Chip (#7090A, Harlan Teklad) bedding is used in cages and colonies are maintained on a standard rodent diet (#2918 Envigo) with unrestricted access to food and water. The University of Kentucky Institutional Animal Care and Use Committee granted approval for all animal studies conducted.

### Body composition analyses

Body composition analysis was performed on conscious mice using an EchoMRI™ 100H instrument at the University of Kentucky’s Energy Balance and Body Composition Core. Measurements of fat and lean mass were divided by directly measured total body mass and multiplied by 100 to calculate fat and lean mass as a percent of total body mass.

### Blood collection and plasma lipid analyses

Blood samples were obtained from mice by retro-orbital bleeding. Blood was collected into 250 µL heparin-coated glass capillary tubes. Plasma was separated from blood by centrifugation at 1,250 x g for 10 minutes at 4°C. Plasma lipid concentrations were measured using colorimetric enzymatic assays, Cholesterol-E (#999-02601, FujiFilm) and L-Type Triglyceride M (#994-02891 and #990-02991, FujiFilm).

### Oral oil gavage study

To measure postprandial plasma triglycerides after an oral fat load, mice were maintained on time-restricted feeding with 8-hour access to food during the day for 4 days prior to the oral gavage experiment. On the day of the experiment, mice received an oral gavage of vegetable oil (10 µL per gram of body weight) using a blunt ball-tipped syringe. Plasma was obtained at 0, 1, 2, and 4 hours post-gavage and triglyceride levels were measured by enzymatic assay.

### Plasma lipase activity

A heparin study was used to determine plasma lipoprotein lipase activity. Blood was collected via retro-orbital bleeding with 50 µL heparinized capillary tubes and plasma separated by centrifugation at 1,250 x g for 10 minutes at 4°C. Pre-heparin plasma was collected at baseline and post-heparin plasma collected 10 minutes after retro-orbital injection of 300 units heparin per kg body weight. The plasma was then used in a lipase activity assay carried out at 37°C in a black 96-well plate in the presence of assay buffer (0.15 M NaCl, 20 mM Tris-HCl, 1.5% BSA, pH 8.0) and lipase substrate (substrate prepared in 0.05% Zwittergent) for a total volume of 100 µL. Assay was read every 2.5 minutes for a total time course of 10 minutes.

### Sucrose polybehenate study

Dietary fat absorption studies were performed using the non-absorbable fatty acid tracer sucrose polybehenate as previously described ^34^. Four-month-old mice were given high-fat diet containing 45% kcal from fat (Research Diets, Cat# D12451) for two weeks. On day 14, this diet was switched to an identical diet except with 5% sucrose polybehenate (SP) added (Research Diets, Cat# D23011202). After three days on SP diet, mice were moved to individual cages with wire mesh bottoms and paper towels. Fecal pellets were collected and used to quantitatively assess the fat absorption efficiency in these mice. The ratios of behenic acid to total fatty acids in the food and in the feces, as determined by gas chromatography of fatty acid methyl esters, were used to assess the % absorption of dietary fatty acids.

### Electron microscopy imaging of intestinal tissue

Electron microscopy of small intestine was performed as described previously ^20^. Mice were euthanized by cervical dislocation and a segment of duodenal small intestine tissue was collected and immediately immersed in Karnovsky’s fixative. The tissue was subsequently fragmented into cubes measuring 1mm^3^ or smaller. The sample was then subjected to post-fixation with osmium tetroxide, dehydration, and finally embedded in Epon for sectioning. The imaging of sections on uncoated grids was performed using a JEOL JEM 1200EXII transmission electron microscope.

### Intestinal protein analysis

During the harvest process, small intestine from mice was first flushed with PBS containing 2x halt^TM^ protease inhibitor (Thermo, #78442). The proximal small intestine was divided into sections and frozen rapidly in liquid nitrogen. The frozen tissue was then stored at −80°C until protein lysates were prepared. Approximately 25 mg of intestinal tissue, between 10-11.5 cm away from the pyloric sphincter for each mouse, was subjected to lysis using RIPA lysis buffer (Thermo, #89901) supplemented with 2x halt^TM^ protease inhibitor (Thermo, #78442). Tissue homogenization was carried out in red Rino tubes (Next Advance) utilizing a Bullet Blender (Next Advance, model #BB24-AU) operating at level-8 speed for a duration of 4 minutes. The lysate underwent centrifugation at 10,000 x g for 10 minutes at 4°C, resulting in the collection of the supernatant. The quantification of total protein was conducted using the Bicinchoninic acid (BCA) protein assay. 10 µg of total protein from intestinal was used for Western blotting. The specific antibodies used for this study were LC3b (R&D Biosystems, #MAB85582), and Vinculin (Novus Biologicals, #NB600-1293), which served as the normalization control.

### Intestinal lipid staining and immunofluorescence

A 5 cm long segment of proximal small intestine, beginning 2 cm distal to the pyloric sphincter, was used to make intestinal swiss rolls. The rolls were fixed in 10% formalin overnight at room temperature and then transferred to 30% sucrose in PBS for 24 h at 4°C prior to being embedded in OCT compound. Embedded rolls were then stored at −80°C until they were sectioned on a cryostat. Sections of 10 μm thickness were cut and mounted on slides which were then used for Oil Red O staining or immunofluorescence experiments. Oil Red O stained intestinal tissue sections were imaged using Zeiss Axioscan Z7 under 20x brightfield settings. Fluorescence images for LC3 immunostained intestinal tissues were obtained using a Nikon A1R confocal microscope. The specific antibodies used for immunofluorescence study were LC3b (R&D Biosystems, #MAB85582), and Alexa fluor^TM^ 568 donkey anti-rabbit IgG (Invitrogen, #A10042).

### Intestinal gene expression by RNA transcriptomics and quantitative PCR

Small intestinal sections, between 8.5-10 cm distal to the pyloric sphincter, were cut into multiple pieces and promptly transferred into RNA Later solution (Invitrogen, #AM7021). These tissues were then stored at a temperature of 4°C prior to proceeding with RNA extraction using the RNAqueous^TM^-4PCR kit (Invitrogen, #AM1914). RNA integrity was analyzed using a TapeStation 4150 instrument (Agilent) and total RNA tape kits. Total RNA was used for RNA sequencing analysis performed by Novogene Co.

### Gut microbiome analysis

For gut microbiome analysis, fresh fecal pellets were collected from individually housed mice (n=5/genotype/sex) in wire-bottom cages. Two fecal pellets per mouse were collected using sterile toothpicks and immediately placed into the sample collection tubes (Transnetyx Microbiome) containing DNA stabilization buffer before being shipped to Transnetyx (Cordova, TN, USA) for DNA extraction, library preparation, and sequencing. After DNA extraction and quality control, the libraries were sequenced at a depth of 2 million 2×150 bp read pairs using the shotgun sequencing technique, which allows for species and strain level taxonomic precision. The sequencing results were analyzed using the One Codex program and compared to a database of over 127K complete microbial reference genomes.

### Mitochondrial energetics measurements

Mitochondrial DNA content was measured as previously described ^35^. Briefly, total (mitochondrial and nuclear) DNA from liver was isolated by phenol/chloroform/isoamyl alcohol extraction. Both mitochondrial (Dloop) and nuclear (Tert) DNA region were amplified by real-time qPCR with 30 ng of DNA (Bio-Rad CFX Connect with Bio-Rad iTaq supermix). The mitochondrial content was expressed as the Dloop/Tert ratio using Standard Quality values. Respirometry in frozen samples was performed on frozen liver biopsies as previously described ^36^. Briefly, liver pieces were homogenized in 500 µl MAS buffer (without BSA), spun at 1000g to remove debris, and protein quantified by Bradford (Bio-Rad, Protein Assay Dye Reagent Concentrate, #5000006). The oxygen consumption rate was measured for complexes I (1 mM NADH), II (5 mM succinate), and IV (0.5 mM TMPD) in presence of 10 µg/ml cytochrome c in MAS (with 0.2% BSA) in a Seahorse Bioscience XF96 instrument. Values were normalized per µg protein.

### Statistical analyses

Unpaired Student’s t-tests were used for direct two-groups comparisons. ANOVA was used for analyses comparing more than two groups, with post-hoc adjustments made according to the methods stated in the figure captions for each experiment. GraphPad Prism was used to perform all statistical analyses. P values < 0.05 were regarded as statistically significant for all experiments.

## Notes

### Competing Interest Statement

The authors have declared no competing interest.

### Summary of Updates

Minor edits to the main text and addition of one panel to Figure 6.

