## Supplemental figures for "Impaired Chylomicron Secretion Results in Reduced Body Fat and Increased Intestinal Fatty Acid Oxidation by Activation of Autophagy"

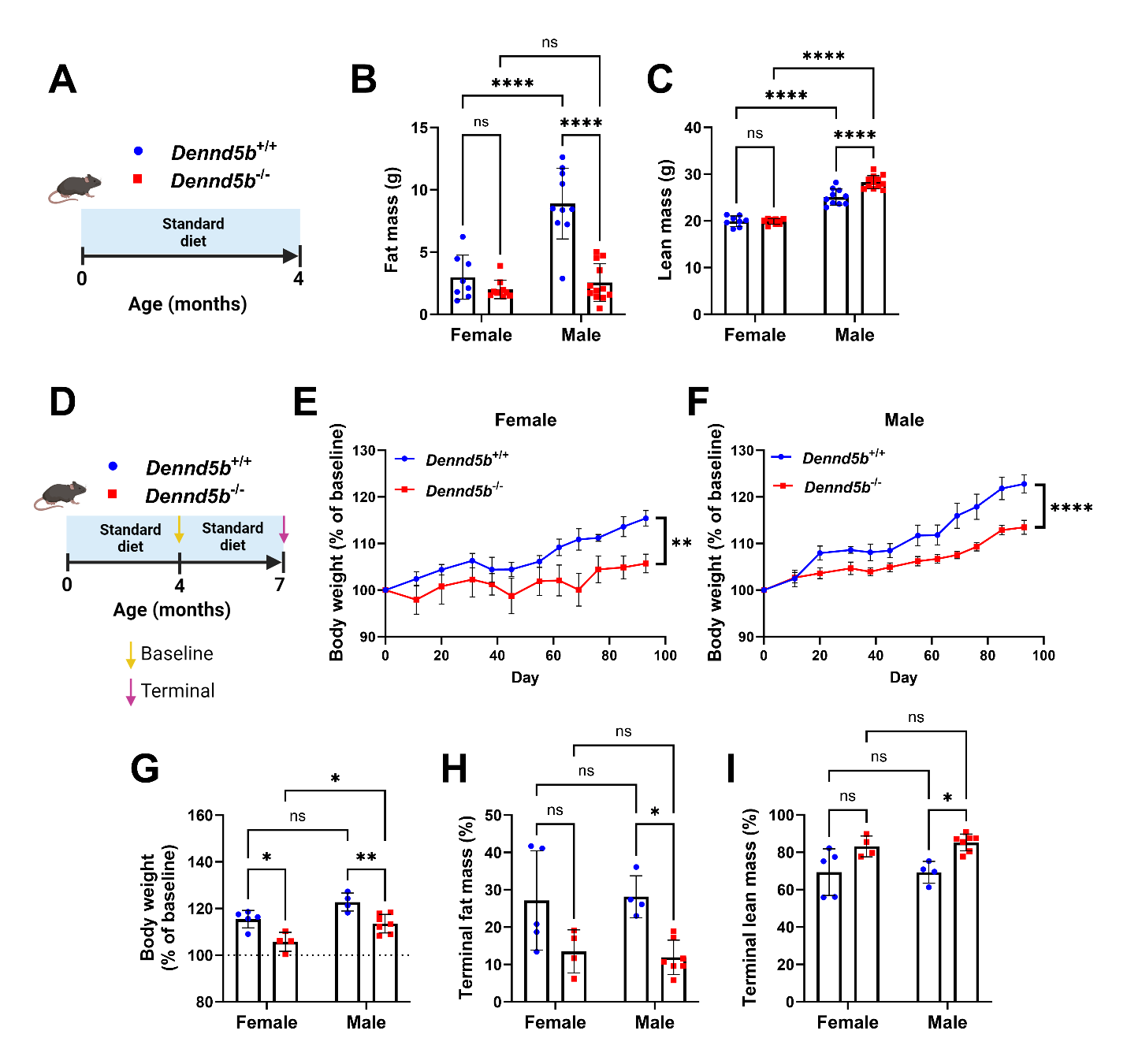
**Supplementary Figure 1. *Dennd5b* is associated with body composition in mice.** (A) Schematic of study design to evaluate body composition in 4-month-old Dennd5b^+/+^ and Dennd5b^-/-^ female and male mice fed standard chow diet (SD). (B, C) Echo MRI body composition of 4-month-old SD fed Dennd5b^+/+^ and Dennd5b^-/-^ mice. n=8-12/group. Data are mean +/- standard error. Comparisons by Two-way ANOVA with Tukey correction for multiple comparisons. (D) Schematic of study design to determine the impact of aging on weight gain in Dennd5b^+/+^ and Dennd5b^-/-^ mice maintained on SD from 4 to 7 months of age. (E-F) Changes in body weight of Dennd5b^+/+^ and Dennd5b^-/-^ female (E) and male (F) mice during 3-months feeding SD. Body weight was measured every 10 days. (G) Percent body weight change on standard diet over a 3-month period (4 to 7 months old). (H-I) Body composition analysis of Dennd5b^+/+^ and Dennd5b^-/-^ mice at the Terminal time point (7-months of age). n=4-7/group. Data are mean +/- standard error. Comparison by Two-way ANOVA with Sidak’s multiple comparisons test. For all statistical comparisons: *p<0.05, **p<0.01, ****p<0.0001.

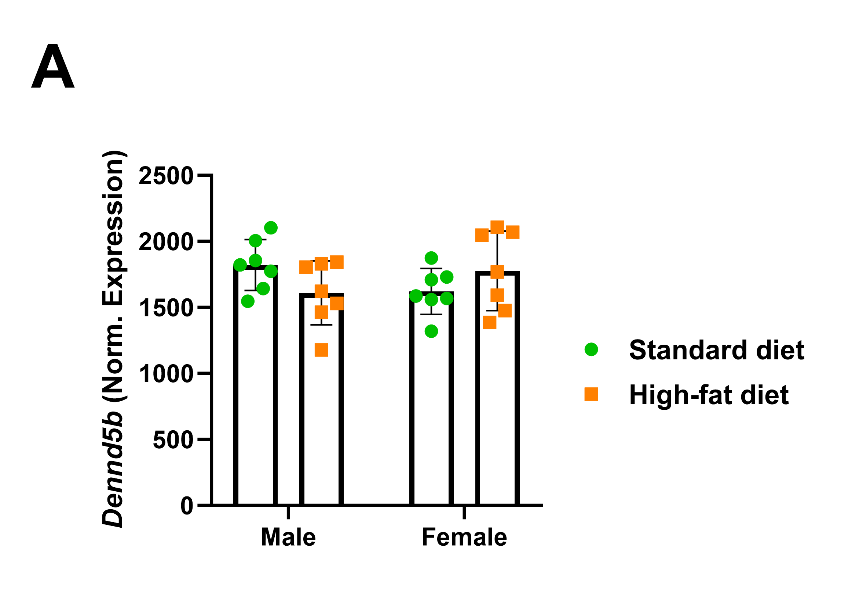
**Supplementary Figure 2. High-fat diet feeding does not impact duodenal *Dennd5b* mRNA expression levels in wild type mice.** (A) Dennd5b mRNA abundance in duodenal tissue from 4-month-old *Dennd5b^+/+^* mice maintained on standard chow diet or high-fat diet (1 month). n=7/group. Data are mean +/- standard error. Comparison by Two-way ANOVA with Sidak’s multiple comparisons test.

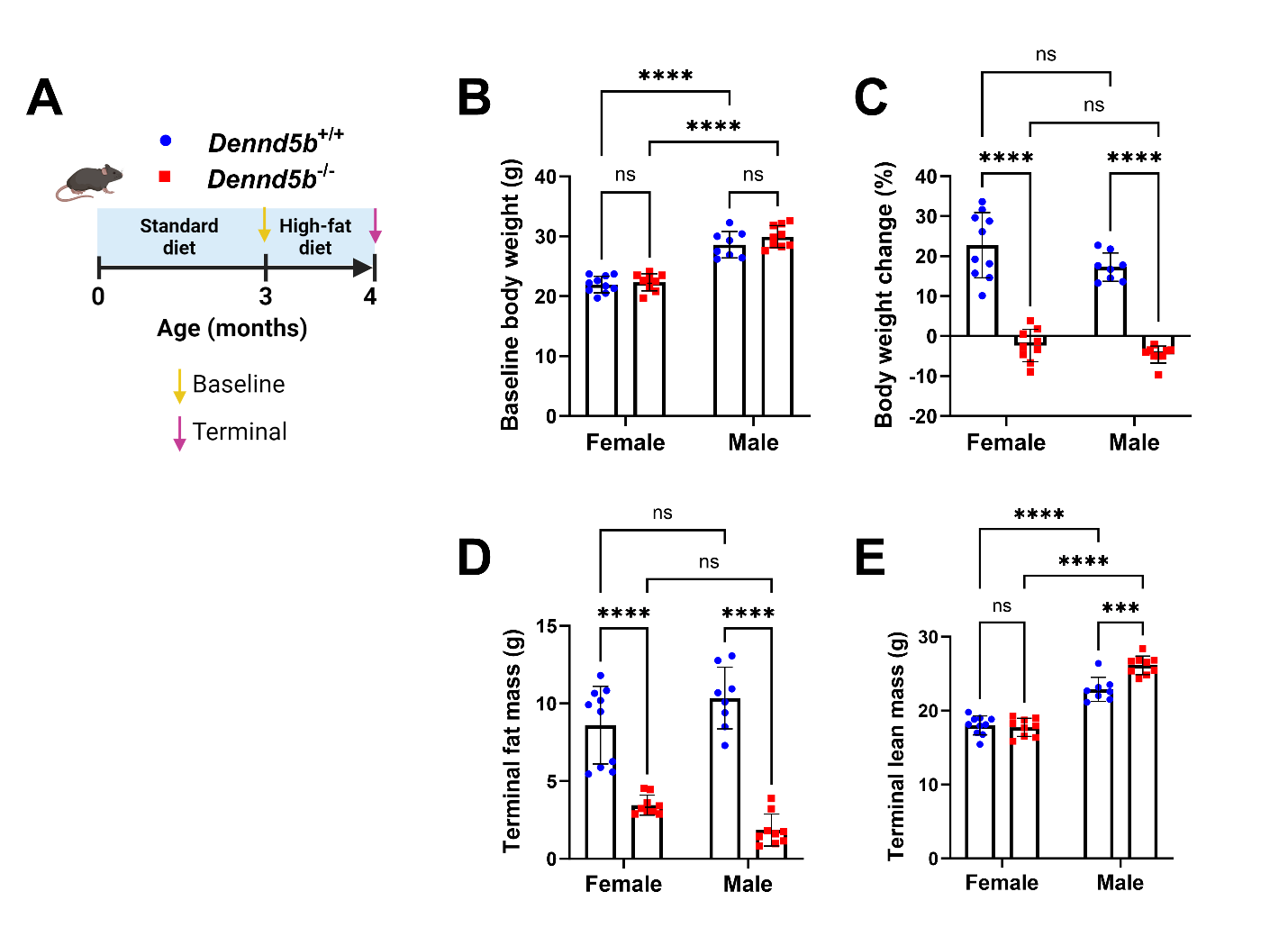
**Supplementary Figure 3. Dennd5b^-/-^ increases lean mass in high-fat diet fed male mice.** (A) Schematic of study design to evaluate body composition in 4-month-old Dennd5b^+/+^ and Dennd5b^-/-^ female and male mice fed high-fat diet. (B) Baseline body weights. (C) Percent change in body weight after one month of high-fat diet feeding. (D-E) Fat and lean body mass after 1 month of high-fat diet feeding. n=8-10/group. Data are mean +/- standard error. Comparison by Two-way ANOVA with Tukey correction for multiple comparisons. For all statistical comparisons: ***p<0.001, ****p<0.0001.

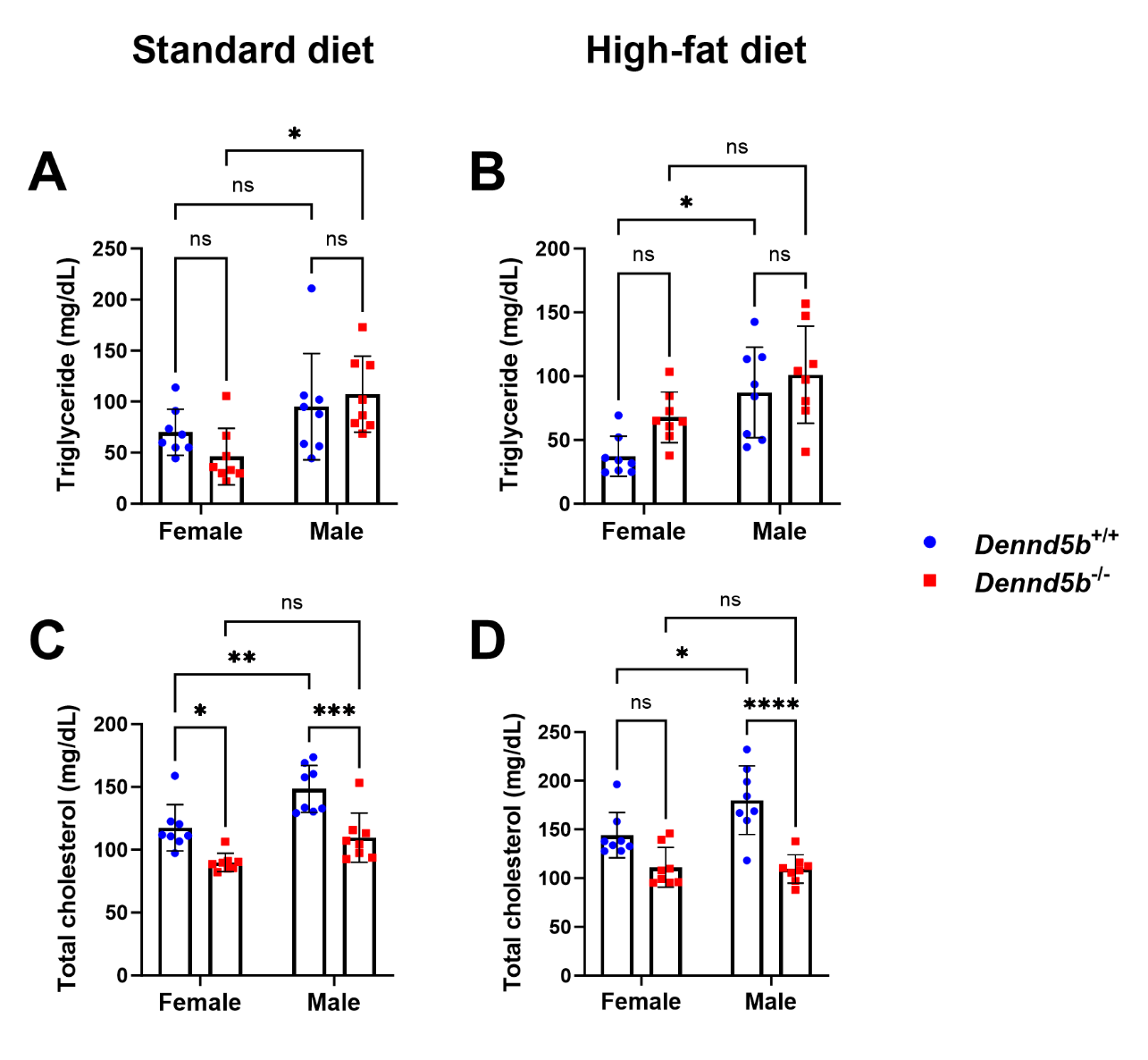

**Supplementary Figure 4. *Dennd5b^-/-^* mice have reduced plasma cholesterol.** Dennd5b^+/+^ and Dennd5b^-/-^ mice were maintained on standard diet until 3 months of age, then continued on standard chow diet or switched to high-fat diet for 1 month. Plasma lipids were measured in 4-month-old mice. (A) Plasma triglyceride concentrations of mice fed standard chow diet. (B) Plasma triglyceride concentrations of mice fed high-fat diet. (C) Plasma total cholesterol concentrations of mice fed standard chow diet. (D) Plasma total cholesterol concentrations of mice fed high-fat diet. Data are mean +/- standard error (n=8/group). Comparison by Two-way ANOVA with Sidak’s multiple comparisons test. For all statistical comparisons: *p<0.05, **p<0.01, ***p<0.001, ****p<0.0001.

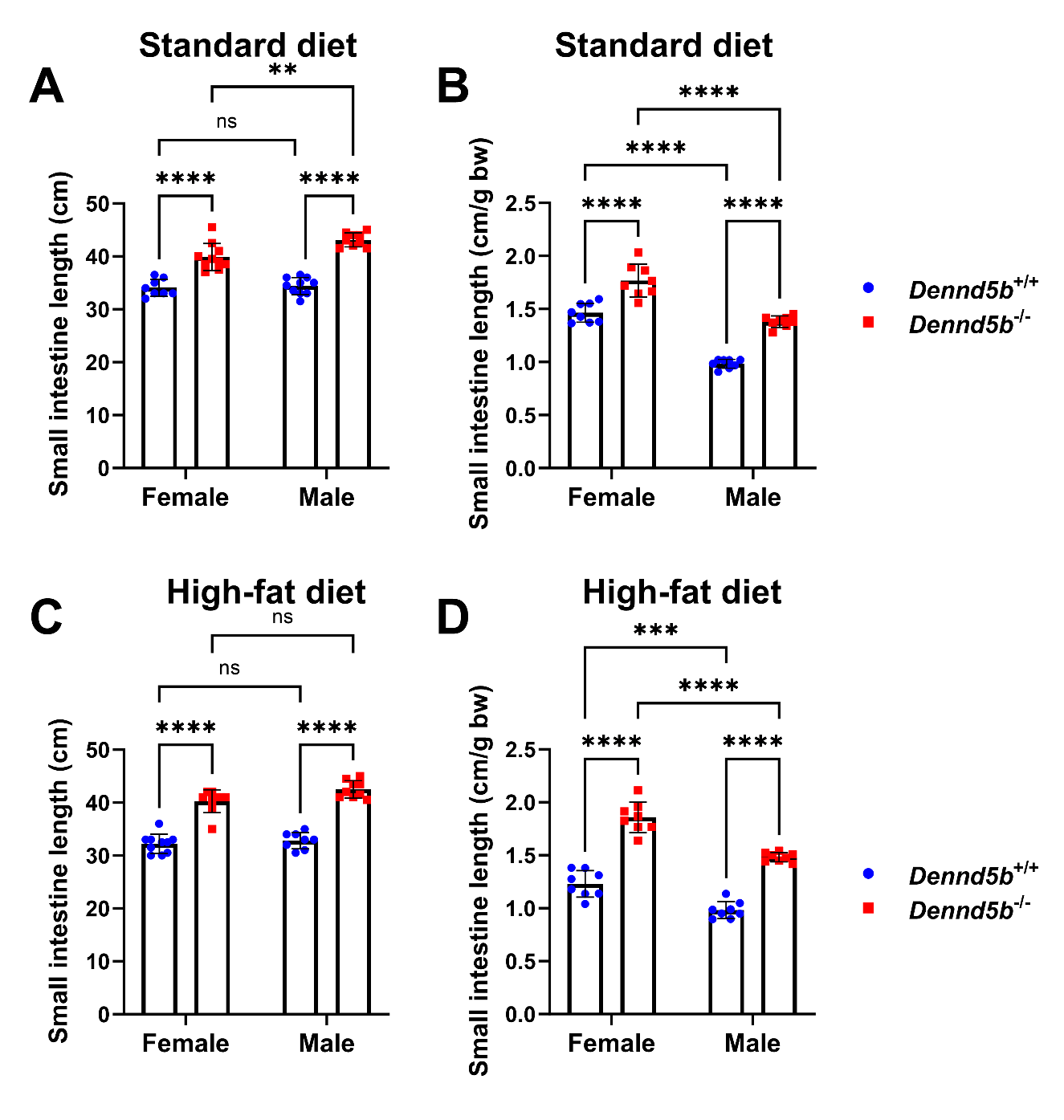

**Supplementary Figure 5. *Dennd5b* deficient mice have longer small intestine.** Analysis of total small intestinal length, not adjusted for body weight, from wild type and *Dennd5b^-/-^* mice maintained standard chow diet (A,B) or high-fat diet (C,D). Data are displayed as total small intestine length (A,C) and the same value adjusted for bodyweight (B,D). Data are mean +/- standard error (n=8/group). Comparison by Two-way ANOVA with Sidak’s multiple comparisons test. For all statistical comparisons: **p<0.01, ***p<0.001, ****p<0.0001.

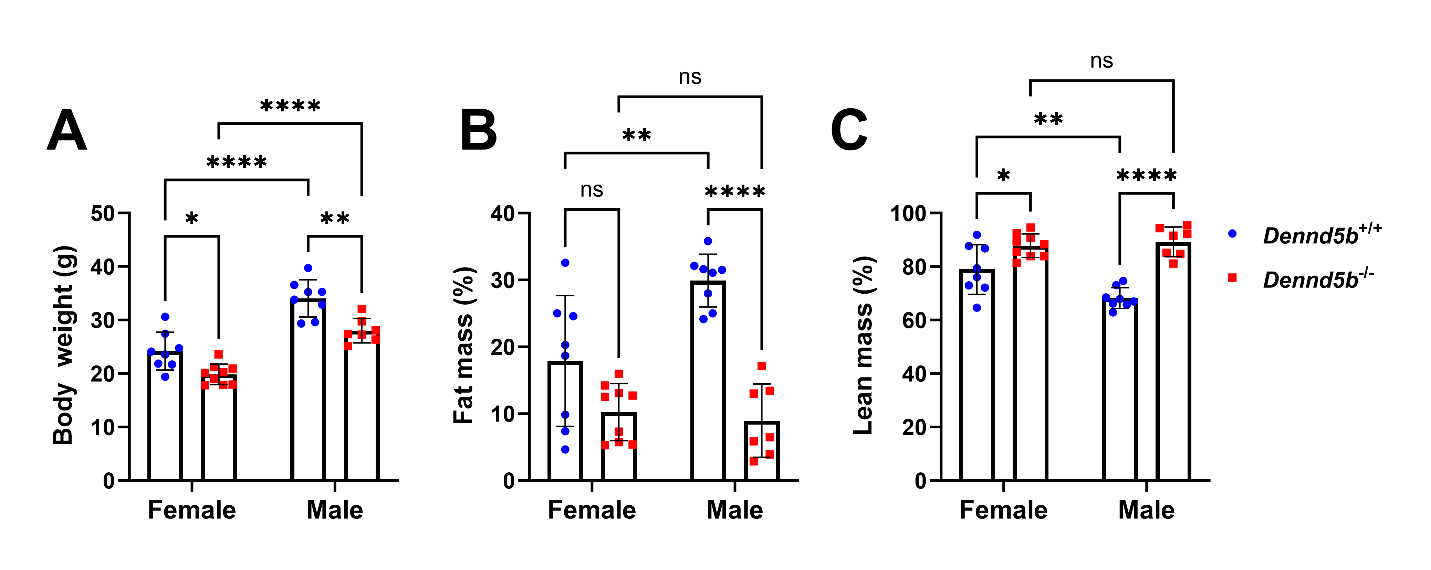

**Supplementary Figure 6. Body composition of mice from the sucrose polybehenate study.** (A) Body weights and (B-C) body composition analyses of Dennd5b^+/+^ and *Dennd5b^-/-^* mice after 3-weeks feeding HFD and HFD-SP. Data are mean +/- standard error (n=7-9/group). Comparison by Two-way ANOVA with Sidak’s multiple comparisons test. For all statistical comparisons: *p<0.05, **p<0.01, ****p<0.0001.

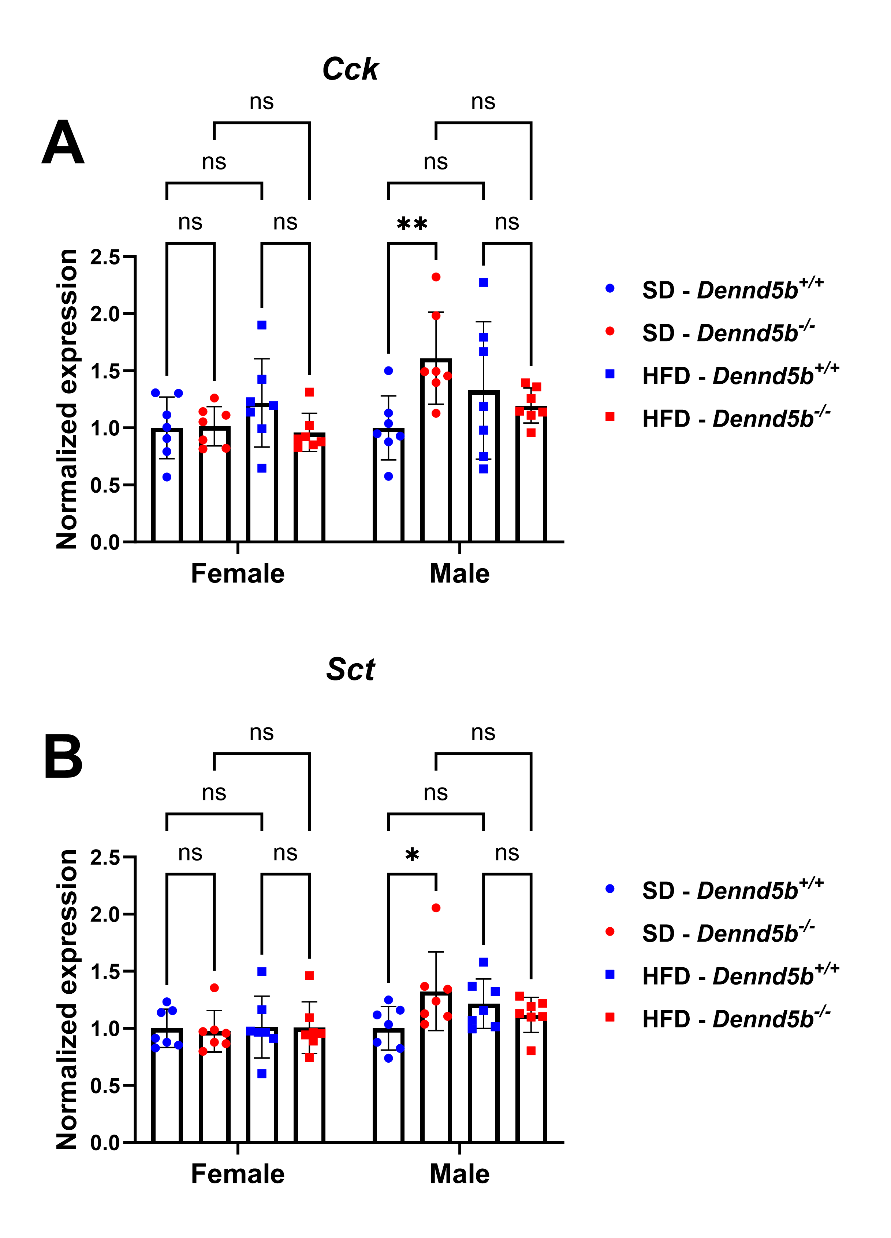
**Supplementary Figure 7. *Dennd5b-*deficiency in mice does not alter expression of endocrine hormones in intestinal tissue**. Gene expression in duodenal intestine tissue from Dennd5b^+/+^ and Dennd5b^-/-^ mice maintained either on standard chow diet (SD) or high-fat diet (HFD) for 1 month. Duodenal mRNA abundance of cholecystokinin (*Cck*) (A) and secretin (*Sct*) (B). Data are mean +/- standard error (n=7/group). Comparison by Two-way ANOVA with Tukey’s multiple comparisons test. For all statistical comparisons: *p<0.05, **p<0.01.

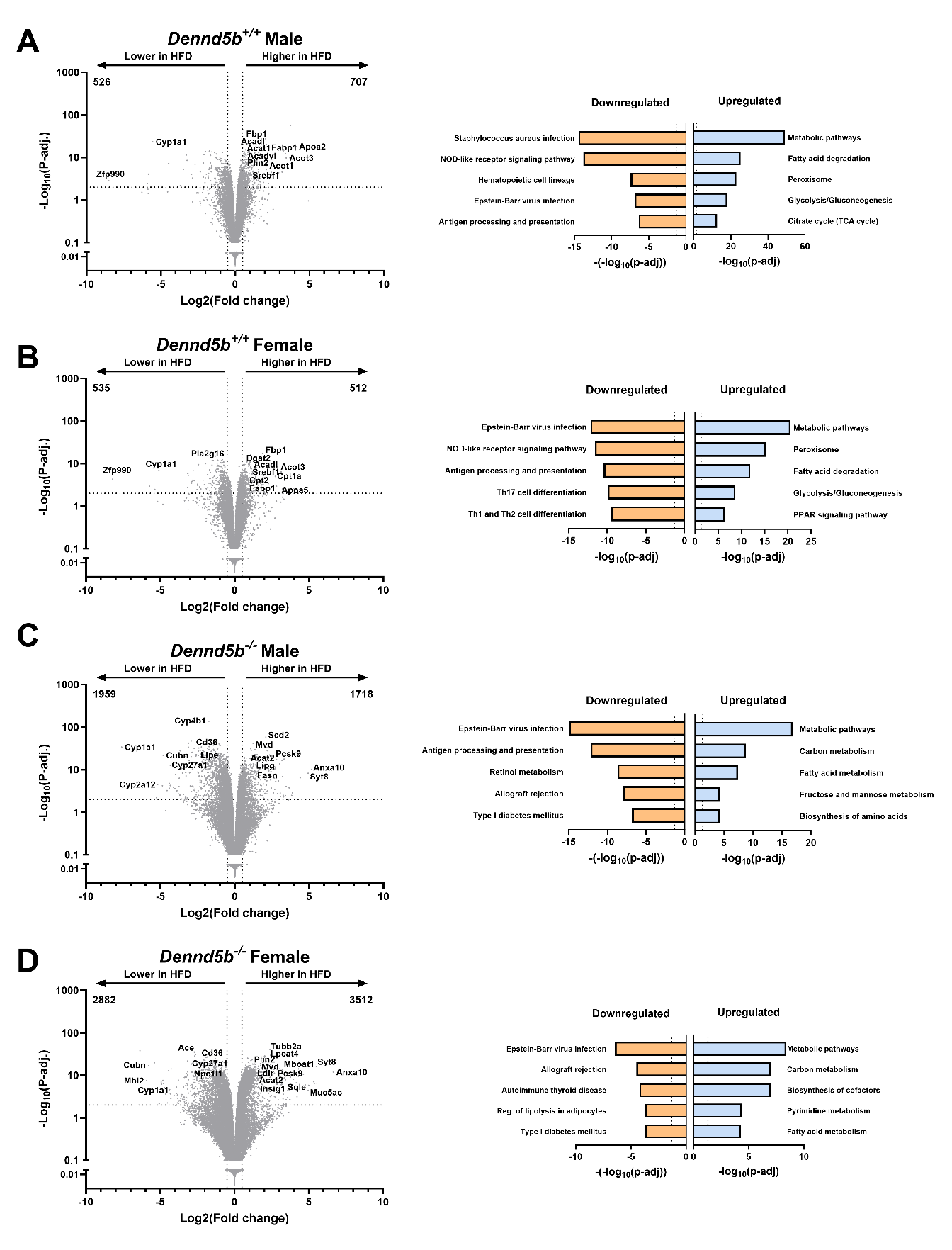

**Supplementary Figure 8. Impact of high-fat diet on intestinal gene expression in wildtype and *Dennd5b* deficient mice.** Volcano plots and Gene Ontology pathway analysis showing RNA abundance and corresponding pathways in duodenal tissue from 4-month-old *Dennd5b^+/+^* and *Dennd5b^-/-^* mice maintained on standard chow diet or high-fat diet for 1 month.

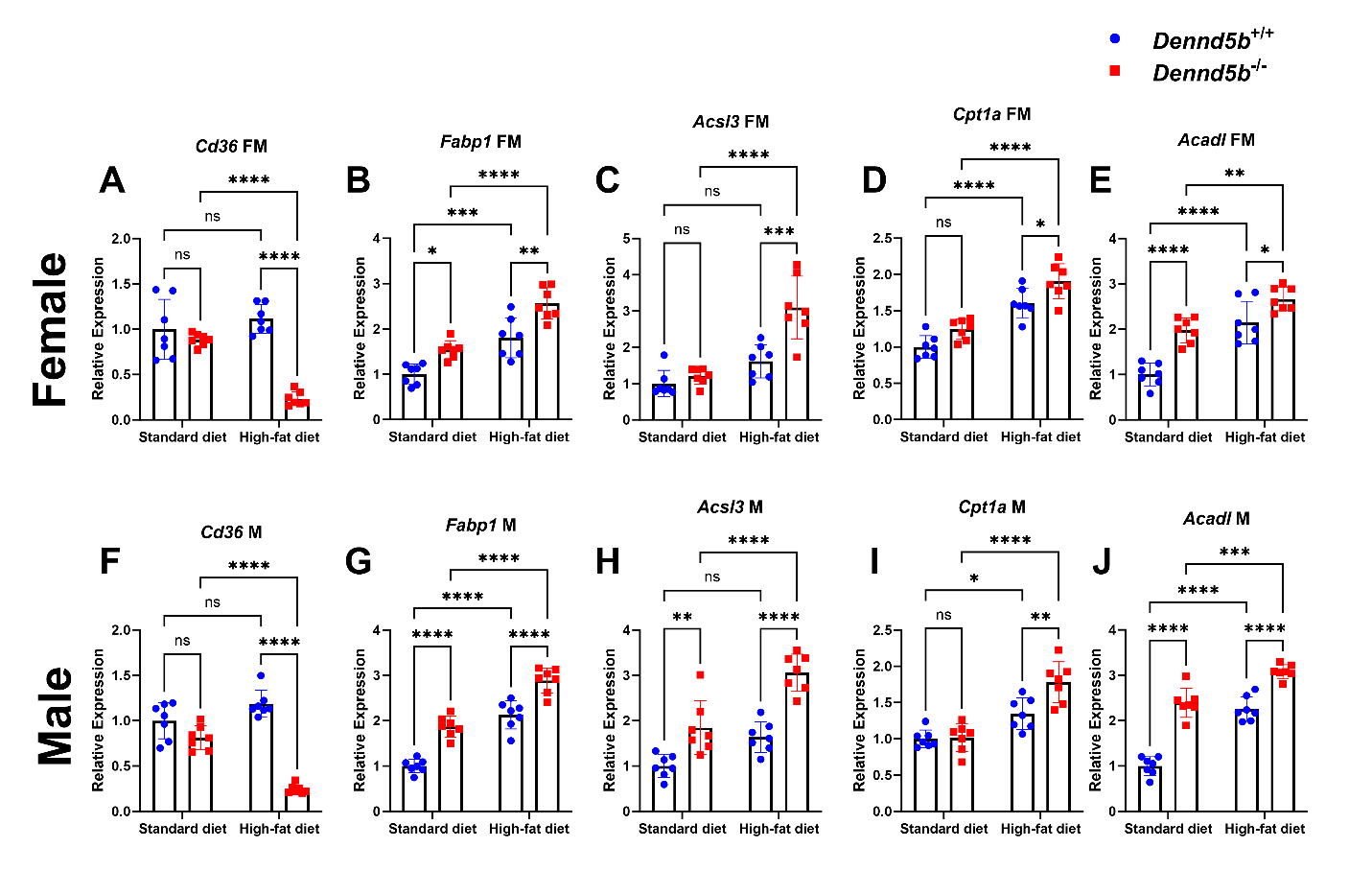
**Supplementary** **Figure 9. *Dennd5b* deficiency impacts mRNA abundance of genes involved in fatty acid beta oxidation.** RNA abundance in duodenal tissue from 4-month-old *Dennd5b^+/+^* and *Dennd5b^-/-^* mice maintained on standard chow diet or high-fat diet for 1 month. (A,F) Duodenal mRNA abundance of *Cd36*, (B,G) *Fabp1*, (C,H) *Acsl3*, (D,I) *Cpt1a*, (E,J) and *Acadl.* Data are mean +/- standard error (n=7/group). Comparison by Two-way ANOVA with Sidak’s multiple comparisons test. For all statistical comparisons: *p<0.05, **p<0.01, ***p<0.001, ****p<0.0001.

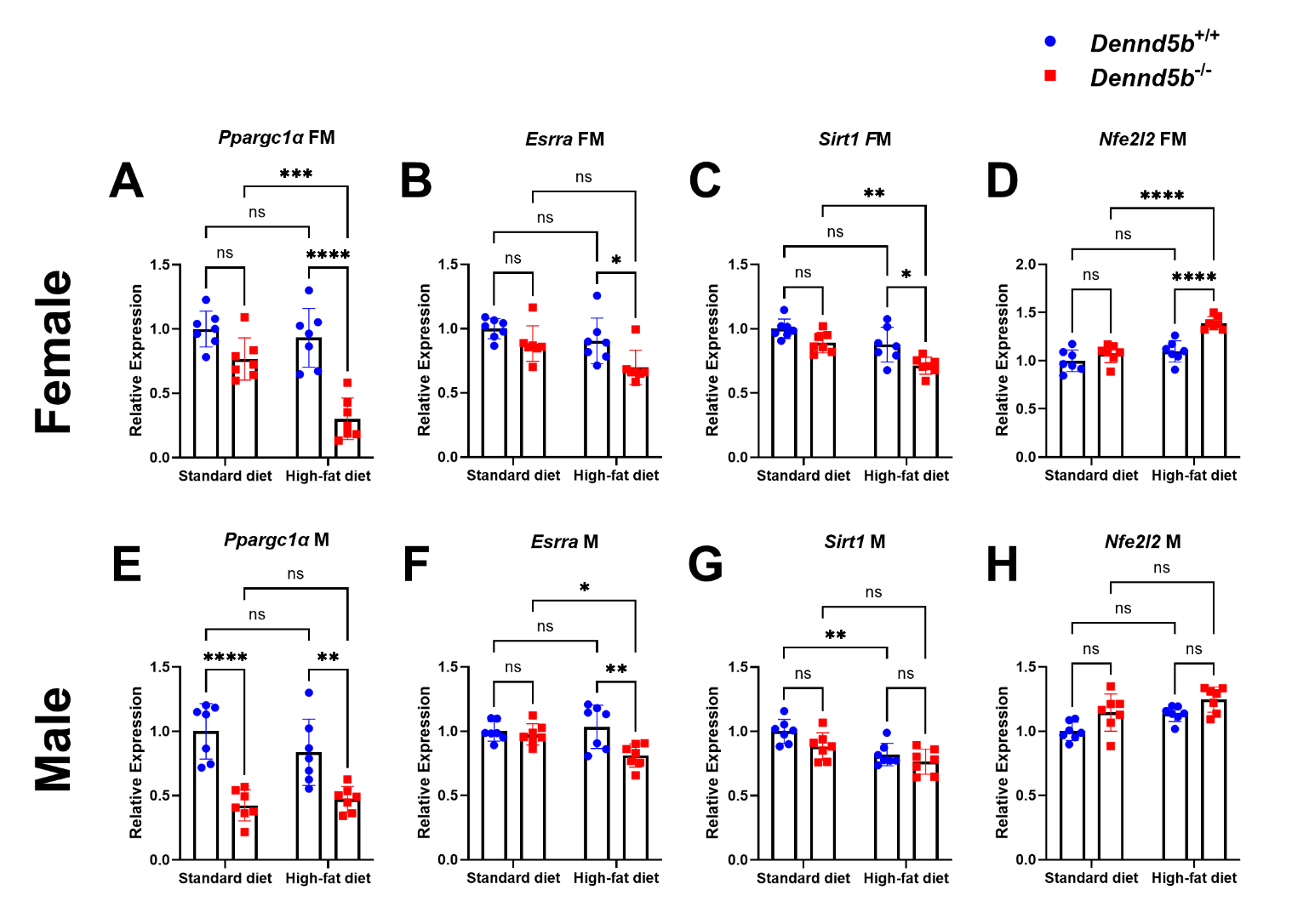
**Supplementary Figure 10. Impact of *Dennd5b* deficiency on mRNA abundance of *Ppargc1α* and other genes involved in regulating cellular mitochondrial content.** *Ppargc1a* mRNA abundance in duodenal tissue from 4-month-old female (A) and male (B) *Dennd5b^+/+^* and *Dennd5b^-/-^* mice maintained on standard chow diet or high-fat diet for 1 month. Data are mean +/- standard error (n=7/group). Comparison by Two-way ANOVA with Tukey’s multiple comparisons test. For all statistical comparisons: **p<0.01, ***p<0.001, ****p<0.0001.

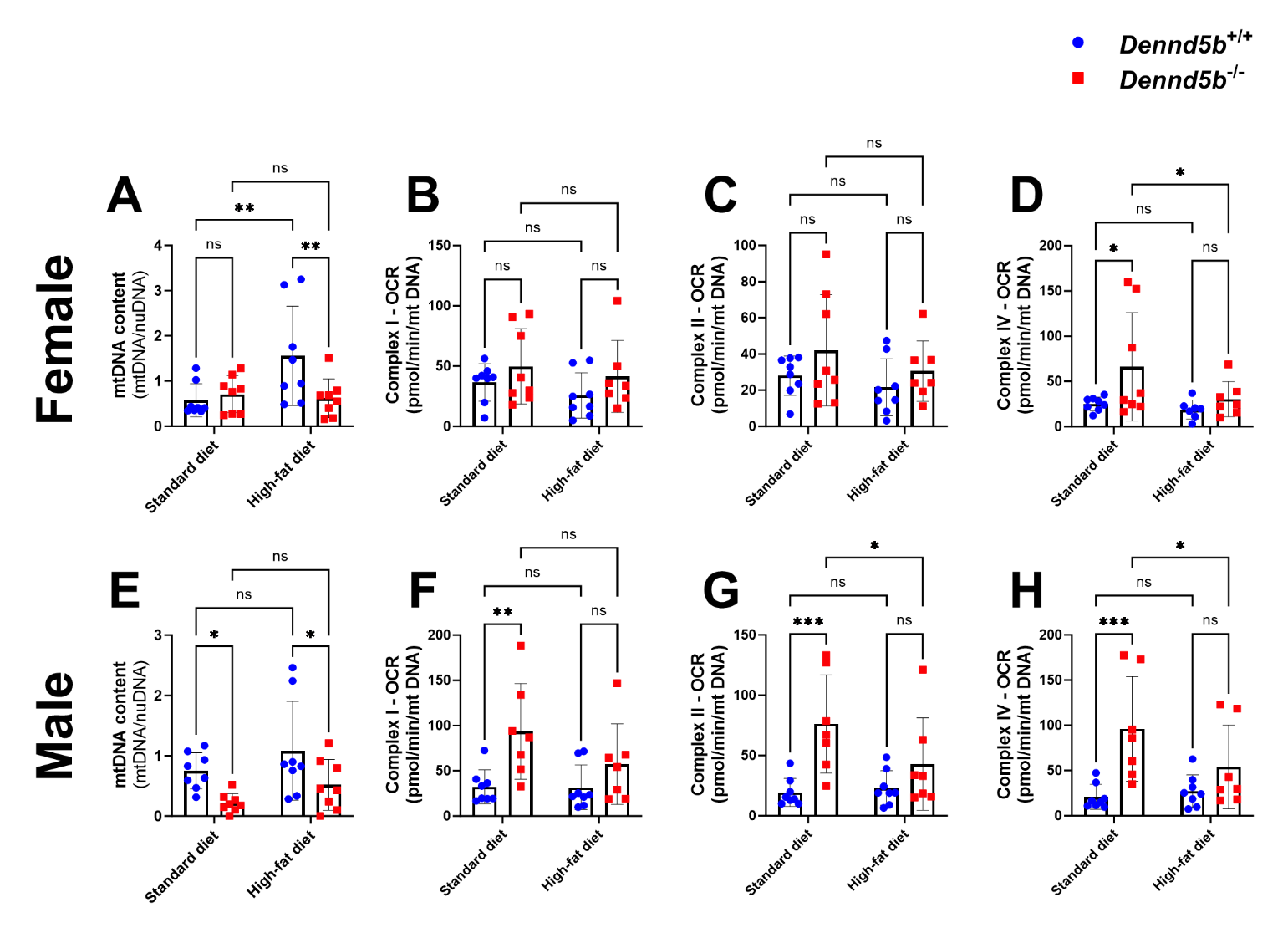

**Supplementary Figure 11. Impact of *Dennd5b* deficiency and high-fat diet on intestine mitochondrial function normalized to mitochondrial DNA content.** Mitochondrial DNA content and oxygen consumption rates measured in duodenal tissue from 4-month-old female and male *Dennd5b^+/+^* and *Dennd5b^-/-^* mice maintained on standard chow diet or high-fat diet for 1 month. Data are mean +/- standard deviation (n=7-8/group). Comparison by Two-way ANOVA with uncorrected Fischer’s LSD multiple comparisons test. For all statistical comparisons: *p<0.05, **p<0.01, ***p<0.001.

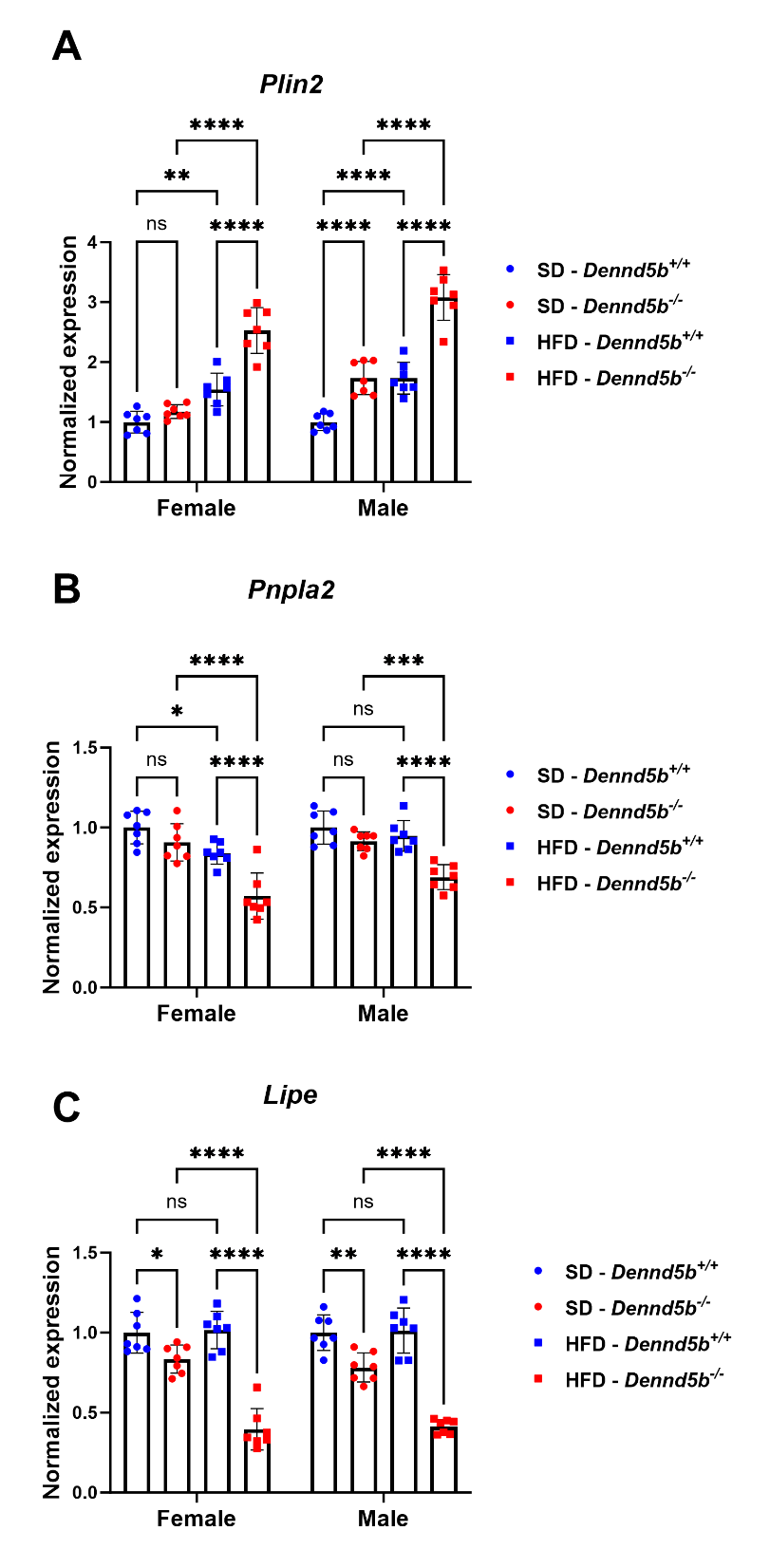

**Supplementary Figure 12. Gene expression suggests increased neutral lipid storage and reduced lipolysis of lipid droplets in *Dennd5b^-/-^* intestine.** (A-C) Duodenal mRNA abundance for lipid droplet surface protein Perilipin 2 (*Plin2*), and triglyceride hydrolysis enzymes adipose triglyceride lipase (*Pnpla2*) and hormone-sensitive lipase (*Lipe*) in female and male *Dennd5b^+/+^* and *Dennd5b^-/-^* intestinal tissue from mice maintained on standard diet (SD) or high-fat diet (HFD) for 1 month. Data are mean +/- standard error (n=7/group). Comparison by Two-way ANOVA with Tukey’s multiple comparisons test. For all statistical comparisons: *p<0.05, **p<0.01, ***p<0.001, ****p<0.0001.

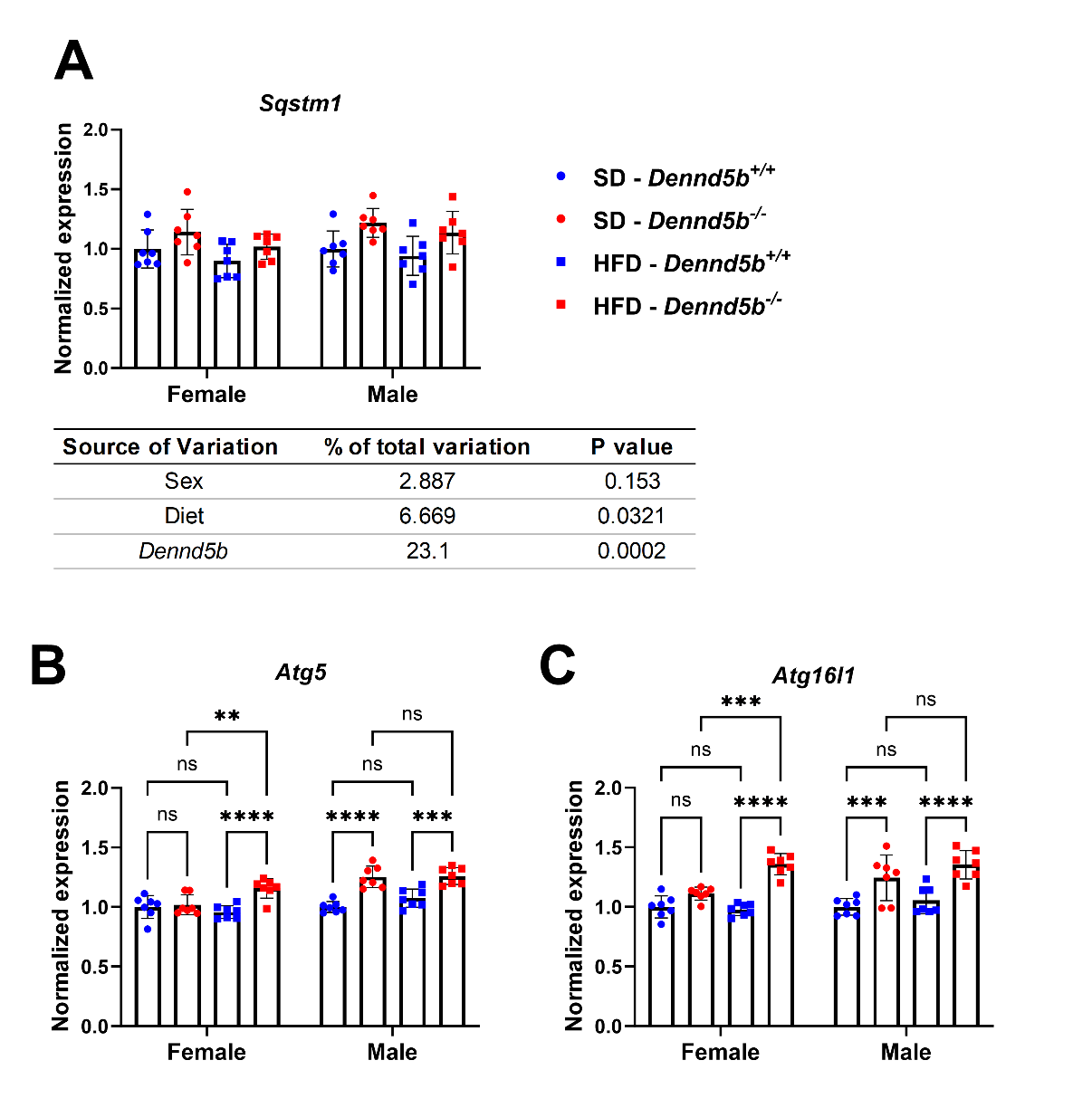

**Supplementary Figure 13. Expression of autophagy related genes in *Dennd5b* deficient mouse intestine.** Duodenal mRNA abundance of Sqstm1 (A), *Atg5* (B), and *Atg16l1* (C) in female and male *Dennd5b^+/+^* and *Dennd5b^-/-^* intestinal tissue from mice maintained on standard diet (SD) or high-fat diet (HFD) for 1 month. Data are mean +/- standard error (n=7/group). Comparisons for A by 3-way ANOVA. Comparisons for B-C by two-way ANOVA with Tukey’s multiple comparisons test. For all statistical comparisons: **p<0.01, ***p<0.001, ****p<0.0001.
